# The centrosomal protein 83 (CEP83) regulates human pluripotent stem cell differentiation towards the kidney lineage

**DOI:** 10.1101/2022.06.20.496810

**Authors:** Fatma Mansour, Christian Hinze, Narasimha Swamy Telugu, Jelena Kresoja, Iman B. Shaheed, Christian Mosimann, Sebastian Diecke, Kai M. Schmidt-Ott

## Abstract

**Background:** During embryonic development, the mesoderm undergoes patterning into diverse lineages including axial, paraxial, and lateral plate mesoderm (LPM). Within the LPM, the so-called intermediate mesoderm (IM) forms kidney and urogenital tract progenitor cells, while remaining LPM forms cardiovascular, hematopoietic, mesothelial and additional progenitor cells. The signals that regulate these early lineage decisions are incompletely understood. Here, we found that the centrosomal protein 83 (CEP83), a centriolar component necessary for primary cilia formation and mutated in pediatric kidney disease, influences the differentiation of human induced pluripotent stem cells (hiPSCs) towards intermediate mesoderm.

**Methods:** We induced inactivating deletions of *CEP83* in hiPSCs and applied a 7 day in vitro protocol of intermediate mesoderm kidney progenitor differentiation, based on timed application of WNT and FGF agonists. We characterized induced mesodermal cell populations using single cell and bulk transcriptomics and tested their ability to form kidney structures in subsequent organoid culture.

**Results:** While hiPSCs with homozygous *CEP83* inactivation were normal regarding morphology and transcriptome, their induced differentiation into IM progenitor cells was perturbed. Mesodermal cells induced after 7 days of monolayer culture of *CEP83*-deficient hiPCS exhibited absent or elongated primary cilia, displayed decreased expression of critical IM genes (*PAX8*, *EYA1*, *HOXB7*) and an aberrant induction of LPM markers (e. g. *FOXF1*, *FOXF2*, *FENDRR*, *HAND1*, *HAND2*). Upon subsequent organoid culture, wildtype cells differentiated to form kidney tubules and glomerular-like structures, whereas *CEP83*-deficient cells failed to generate kidney cell types, instead upregulating cardiomyocyte, vascular, and more general LPM progenitor markers.

**Conclusion:** Our data suggest that *CEP83* regulates the balance of intermediate mesoderm and lateral plate mesoderm formation from human pluripotent stem cells, identifying a potential link between centriolar or ciliary function and mesodermal lineage induction.

## Introduction

During mammalian embryonic development, the mesoderm forms axial, paraxial, and lateral plate domains that harbor precursor cells for distinct organ systems. Forming as a major part of the lateral plate mesoderm (LPM), the intermediate mesoderm (IM) harbors progenitor cells of all kidney epithelial cells^1^, whereas remaining LPM contributes progenitors of various cell types, including cells of the cardiovascular system^2^. The molecular and cellular mechanisms that drive induction of the IM and distinct LPM domains during embryonic development are not fully understood.

The centrosomal protein 83 (CEP83) is a component of distal appendages (DAPs) of centrioles. DAPs are involved in the anchoring of the mother centriole to the cell membrane, an early and critical step in ciliogenesis^3–11^. CEP83 recruits other DAP components to the ciliary base, and loss of CEP83 disrupts ciliogenesis^4^. In radial glial progenitors, removal of CEP83 disrupts DAP assembly, and impairs the anchoring of the centrosome to the apical membrane as well as primary ciliogenesis^5, 10^. Mutations of CEP83 in humans have been associated with infantile nephronophthisis^9^, an early onset kidney disease that results in end stage renal disease before the age of 3 years^12, 13^ and additional organ anomalies^9^. To date, how loss of CEP83 function contributes to aberrant kidney development remains unclear.

Human induced pluripotent stem cells (iPSCs) provide useful tools to study molecular mechanisms of cellular differentiation. Protocols for the induction of kidney organoids from iPSC have been successfully developed^14–19^. The protocol by Takasato *et al*. uses stepwise exposure of iPSC to WNT and FGF agonists in a monolayer culture system for a 7 day period, which results in the induction of cells with a transcriptional phenotype resembling kidney progenitors in the IM^17^. Transfer of these cells to an organoid culture system followed by another series of WNT and FGF signals results in differentiation of 3-dimensional kidney organoids composed of different kidney cells types, including glomerular and tubular cells. Genome editing studies have previously been used to study the effects of genetic defects associated with kidney diseases on kidney differentiation in human iPSC systems ^18, 20–24^. Here, we studied the effect of an induced knockout of *CEP83* in human iPSCs on kidney organoid differentiation. We uncovered a novel role of CEP83 in determining the balance of IM versus LPM differentiation, implicating a centrosomal protein in early mesodermal lineage decisions.

## Concise Methods

### hiPSCs cell line

We used the human iPSC cell line BIHi005-A, which was generated by the Berlin Institute of Health (BIH). The hiPSCs were maintained in 6-well plates (Corning®, 353046) coated with Matrigel (Corning®, 354277) and cultured in Essential 8 medium (E8, A1517001, Gibco-Thermo Fisher Scientific) supplemented with 10µM Y-27632 (Rocki, Wako, 253-00513).

### CRISPR CAS9 Technology to generate CEP83^-/-^ hiPSCs clones

Clustered Regularly Interspaced Short Palindromic Repeats (CRISPR)-Cas9 technology was used to generate *CEP83^-/-^* hiPSCs clones. We designed two CRISPR RNAs (crRNAs) (5’-GGCTGAAGTAGCGGAATTAA-AGG-3’ and 5’-AAGAATACAGGTGCGGCAGT-TGG-3’) using CRISPOR software^25^. The two crRNAs were annealed with trans-activating CRISPR RNA (tracrRNA) to form two guide RNAs (gRNA1 and gRNA2) and then formed a RNP complex by incubating gRNA1 and gRNA2 separately with Alt-R® S.p. Cas9 Nuclease V3 (1 μM concentration, IDT, 1081058).The hiPSCs were transfected with RNP complexes using Neon transfection system (Thermo Fisher Scientific, MPK5000)^26^ and Neon™ transfection 10 μl kit (Thermo Fisher Scientific, MPK10025) according to the manufacturer’s instructions. After 48hrs of transfection, we analyzed the editing efficiency in the pool by PCR genotyping.

For PCR genotyping, we isolated genomic DNA from the pool of transfected cells followed by PCR using Phire™ Tissue Direct PCR Master Mix (Thermo Scientific, F170S) according to the manufacturer’s instructions (**Figure 1B**). After confirming the editing efficiency in the pool, we generated single cell clones by the clonal dilution method. We plated 500 single cells per well of a 6 well plate and picked 24 clones using a picking hood S1 (Max Delbrück Centre Stem Cell Core Facility). Then, clones were screened for homozygous deletions of *CEP83* by PCR using Phire™ Tissue Direct PCR Master Mix. Selected knockout clones were further characterized for *CEP83* loss of function on the DNA, RNA, and protein level. *CEP83^-/-^* clones (*KO1, KO2*, and *KO3*) were registered as (BIHi005-A-71, BIHi005-A-72, and BIHi005-A-73) in the European Human Pluripotent Stem Cell Registry (https://hpscreg.eu).

**Figure 1:**
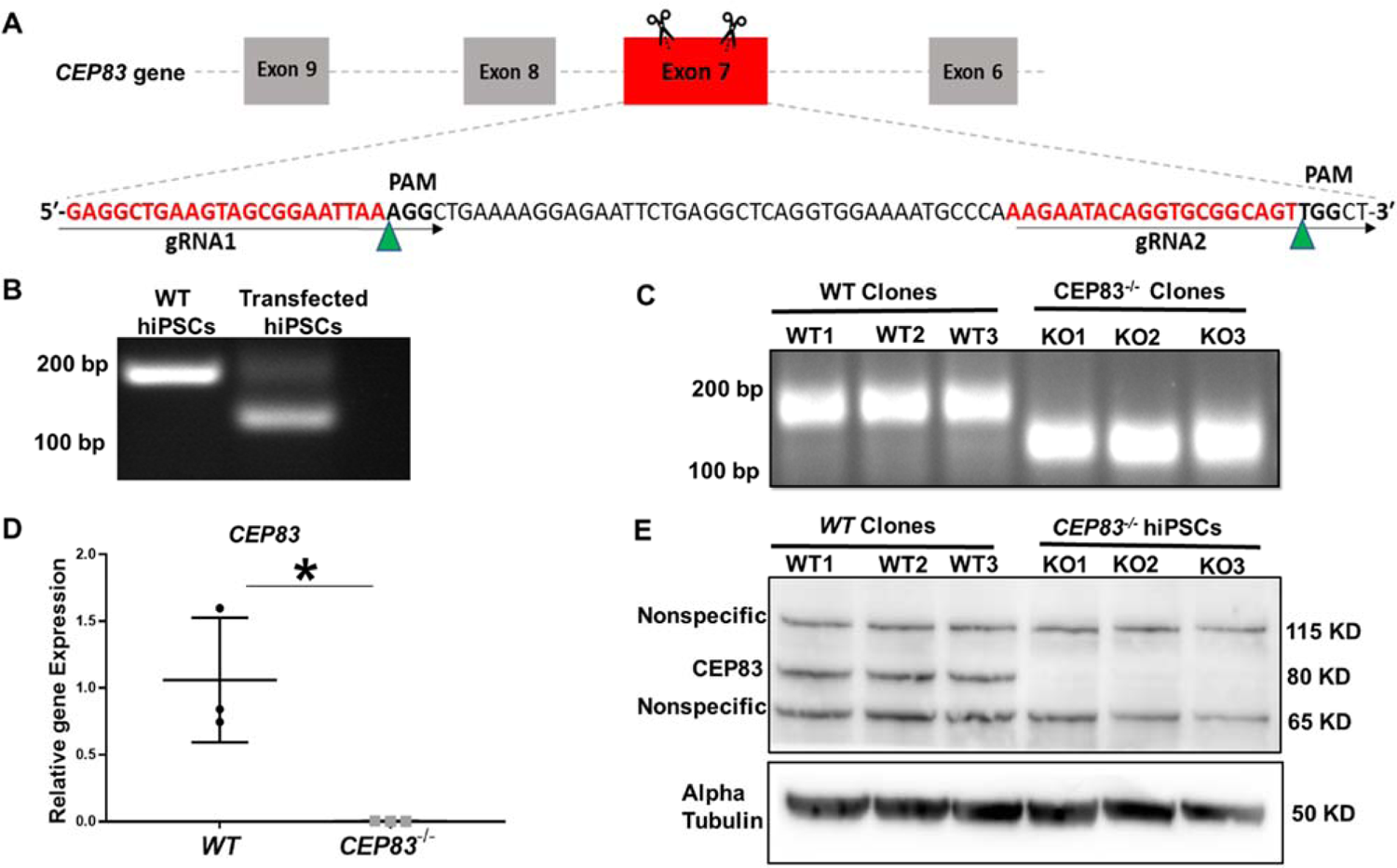
Generation of CEP83-deficient human pluripotent stem cells. (A) Schematic diagram of the experimental approach to induce a deleting mutation in exon 7 of the *CEP83* gene. Two guide RNAs (gRNAs) were designed to induce an approximately 63 bp deletion within exon 7 of the *CEP83* gene after non-homologous end joining. (B) Ribonucleoprotein (RNP) complex containing crispr RNAs (crRNAs), trans-activating crRNA (tracrRNA), and Cas9 endonuclease was transfected into hiPSCs by electroporation. DNA extracted from pooled transfected cells was subjected to PCR targeting the predicted deletion site in the *CEP83* gene. In addition to the 182 bp fragment present in untransfected wildtype (WT) cells, an approximately 120 bp fragment was detected in transfected cells, corresponding to the induced deletion in exon 7. Twenty-four single IPS cell-derived clones from these transfected cells were picked and cultured. (C) Three of these clones (*CEP83*^-/-^ clones *KO1, KO2, KO3*) carried 62-74b bp deletions within *CEP83* exon 7, which led to an induction of premature stop codons or frameshift mutation on both alleles of *CEP83*. Three wildtype clones (*WT1, WT2,* and *WT3*) were used as controls. (D) Quantitative RT-PCR for a fragment corresponding to the deleted region in *CEP83* exon 7 produced a detectable signal in RNA extracts from WT clones but not CEP83^-/-^ clones. (E) Immunoblotting of *WT* and *CEP83^-/-^* clones using a CEP83 antibody targeting the C-terminal region of the protein (see methods for details) indicated a complete loss of the 83 kDa band corresponding to CEP83 protein in the three *KO* clones compared with the three *WT* clones. Data are mean ± SD.*\*P* < 0.05 and *\*\*P* < 0.01 vs. WT. See Figure 1-source data 1-2. See also Figure 1—figure supplements 1–2.

### Single nucleotide polymorphism (SNP)-Karyotype

To assess karyotype integrity, copy number variation (CNV) analysis on the human Illumina OMNI-EXPRESS-8v1.6 BeadChip was used. In brief, genomic DNA was isolated from three *WT* (*WT1, WT2*, and *WT3*) and three *KO* (*KO1*, *KO2* and *KO3*) clones using the DNeasy blood and tissue kit (Qiagen, Valencia, CA, United States), hybridized to the human Illumina OMNI-EXPRESS-8v1.6 BeadChip (Illumina), stained, and scanned using the Illumina iScan system according to a standard protocol^27–29^. The genotyping was initially investigated using the GenomeStudio 1 genotyping module (Illumina). Following that, KaryoStudio 1.3 (Illumina) was used to perform automatic normalization and identify genomic aberrations in detected regions by generating B-allele frequency and smoothed Log R ratio plots. To detect copy number variations (CNVs), the stringency parameters were set to 75 kb (loss), 100 kb (gain), and CN-LOH (loss of heterozygosity). KaryoStudio generates reports and displays chromosome, length, list of cytobands, and genes in CNV-affected regions.

### Differentiation Protocol

We used the protocol of *Takasato* to differentiate the hiPSCs into nephron organoids^17^. Briefly, hiPSCs were cultured firstly in APEL2 medium (Stem Cell Technologies, 05270) supplemented with 5% Protein Free Hybridoma Medium II (PFHMII, GIBCO, 12040077), and 8 μM CHIR99021 (R&D, 4423/10) for 5 days, with medium changes every 2 days. Then, the cells were cultured in APEL2 medium supplemented with 200 ng/ml FGF9 (R&D, 273-F9-025) and 1 μg/ml heparin (Sigma Aldrich, H4784-250MG) for 2 days. On day 7, the cells were washed with 1X Dulbecco’s PBS (DPBS, Thermo Fisher Scientific,14190-250), then trypsinized using trypsin EDTA-0.05% (Thermo Fisher Scientific, 25300-062) at 37 °C for 3 min. The cells were counted and divided to achieve 1×10^6^ cells per organoid and cultured into 3D organoid culture on 0.4-μm-pore polyester membrane of Corning 6-well Transwell cell culture plate (Corning-Sigma Aldrich, CLS3450-24EA). Four to five organoids were seeded on one membrane using a P100 wide-bore tip, and cultured in APEL2 with 5 μM CHIR99021 at 37°C for 1h (CHIR99021 pulse). After the CHIR pulse, we changed the medium to APEL2 medium supplemented with 200 ng/ml FGF9 + 1 μg/ml heparin for 5 days with medium refreshing every 2d. The organoids were then cultured only in APEL2 medium with 1 μg/ml heparin for additional 13 days. The total differentiation time is 25 days (7+18).

### DNA isolation and Polymerase Chain Reaction (PCR)

DNA was isolated from cells using DNeasy Blood & Tissue Kits (Qiagen, 69504). CEP83 primers were designed using Primer3 webtool (Table S1). PCR was done using Phusion high-fidelity DNA polymerase (Biolabs, New England, M0530) according to the manufacturer’s instructions. PCR results were visualized on 1.5% agarose gel using a BioDoc Analyze dark hood and software system (Biometra).

### RNA isolation, RNA Sequencing, and Quantitative PCR (qPCR)

Total RNA was isolated from the cells using RNAasy Mini Kit (QIAGEN, Hilden, Germany, 74104) following the manufacture instructions. The concentration, quality, and integrity of the extracted RNA were evaluated using Nanodrop (Thermo Scientific, Waltham, MA; USA), an Agilent 2100 Bioanalyzer, and the Agilent RNA 6000 Nano kit (Agilent Technologies, 5067-1511). 0.4 μg total RNA was used to obtain a poly A–enriched RNA library by Novogene (Cambridge, United Kingdom). Library concentration was performed using a Qubit fluorometer (HS RNA assay kit, Agilent Technologies). Library size was measured by Agilent 2100 bioanalyzer. The libraries were then subjected to 150-bp paired-end next-generation sequencing (Illumina NovaSeq 6000 S4 flow cells). Mutation visualization was performed using the Integrative Genomic Viewer (IGV) tool^30^. Read counts of the sequenced RNA were normalized to Transcripts Per Million (TPM). The TPM values of the variables were used to plot heatmaps and for principle cell analysis (PCA) based on Pearson correlation, using self-written scripts in R (R Development Core Team (2011)) (version 4.0.4).

RNA was reverse transcribed using the RevertAid First Strand cDNA Synthesis Kit (Thermo Scientific). qPCR was performed using the FastStart Universal SYBR Green Master (Rox) mix (Hoffmann-La Roche) according to the manufacturer’s instructions. Glyceraldehyde-3-Phosphate Dehydrogenase (*GAPDH*) mRNA expression and calculated according to the ΔΔCt method. All primer pairs were designed using Primer3, purchased at BioTeZ (Berlin, Germany), and sequences are shown in (Table S1).

### Single cell RNA sequencing (scRNA-seq) Cells isolation and preparation

Differentiated cells at day 7 were washed twice with 1X DPBS, dissociated with Accumax solution, and resuspended in 1X DPBS. Then, cells were filtered, counted, and checked for viability.

### Library preparation and single cell sequencing

Single cell 3’ RNA sequencing was performed using the 10x Genomics toolkit version v3.1^31^ according to the manufacturer’s instructions aiming for 10000 cells. Obtained libraries were sequenced on Illumina NextSeq 500 sequencers.

### Single-Cell Sequencing Data analysis and Clustering

After sequencing and demultiplexing, fastq files were analyzed using Cellranger version 3.0.2. Gene expression matrices were then imported in R and Seurat objects were created using the Seurat R package (version 4.0.5)^32^. The gene expression matrices were initially filtered by applying lower and upper cut-offs for the number of detected genes (500 and 6000, respectively). The filtered data were then log normalized and scaled according to the number of unique molecular identifiers (UMIs). The normalized and scaled data derived from the four samples were then merged into one Seurat object. Clustering was performed using the first 20 principal components. We used the Seurat FindAllMarkers function to extract marker gene lists that differentiate between clusters with log fold-change threshold ±0.25 using only positive marker expressed in a minimum of 25 % of cells. Principal component analysis (PCA) was done using the first 20 principle components in R using the following libraries factoextra, FactoMineR, and ggplot2.

### Protein extraction and Immunoblotting

Proteins were extracted from hiPSCs using radioimmunoprecipitation assay (RIPA) buffer (Sigma-Aldrich, R0278) as described in details in supplementary data. 30 µg protein in RIPA buffer were mixed with 1x reducing (10% b-mercaptoethanol) NuPAGE loading buffer (Life Technologies, Carlsbad, CA), loaded on a precast polyacrylamide NuPage 4-12% Bis-Tris protein gel (Invitrogen, Carlsbad, CA, USA), and blotted on 0.45 µm pore size Immobilon-P Polyvinylidene difluoride (PVDF) membrane (EMD Millipore, Billerica, MA; USA). The membrane was blocked in 5% bovine serum albumin for 1 h at room temperature and incubated overnight at 4°C with primary antibodies: Anti-CEP83 produced in rabbit (1:500, Sigma-Aldrich) and Anti-α-Tubulin produced in mouse (1:500, Sigma-Aldrich, T9026). Then, the membrane was incubated for 1 h at room temperature with horseradish peroxidase-conjugated secondary antibodies (1:2000, Sigma-Aldrich, Saint Louis, MO, USA). Chemiluminescent reagent (Super Signal–West Pico; Thermo Scientific, Waltham, MA; USA) was used to detect the proteins. The spectra^TM^ Multicolor Broad Range Protein Ladder (Thermo Fisher Scientific, USA) was used to evaluate the molecular weight of corresponding protein bands.

### Histology and Immunofluorescence (IF) staining

Cells at different time points were checked regularly under confocal microscope (Leica DMI 6000 CEL) for differentiation progress. Quantitative analysis of nephron-like structure formation within each organoid (D25) were done on tile scanning images of each organoid by estimating the percentage of the organoid area composed of nephron-like structures using 13 WT and 9 KO organoids. Organoids were fixed in BD Cytofix buffer (554655, BD Biosciences) for 1 hour on ice. Then organoids were gradually dehydrated in increasing ethanol concentrations, cleared in xylene, and embedded in paraffin. Organoids were cut into 3.5 µm-thick sections. The sections were deparaffinized, dehydrated, and stained in hematoxylin (Sigma-Aldrich, Saint Louis, MO) for 3 minutes and in 1% eosin (Sigma-Aldrich) for 2 minutes. For immunostaining, organoids were fixed with BD Cytofix, permeabilized with BD Perm/Wash (554723, BD Biosciences), and blocked with blocking solution (1% BSA + 0.3% triton-X-100 in 1X DPBS) for 2 h. Cells were incubated overnight at 4°C with primary antibodies (table S2), then incubated with fluorescence-labeled secondary antibodies with 1:500 dilution including Cy3, Cy5, Alexa488, and Alexa647 (Jackson ImmunoResearch, Newmarket, UK) and Cy3 Streptavidin (Vector lab, Burlingame, USA) overnight at 4°C. DAPI was then used for nuclear staining (Cell signaling Technology, Danvers, MA, USA) with 1:300000 dilution for 1 hour at RT. Finally, cells were mounted with Dako fluorescent mounting medium (Agilent Technologies). Images were taken using a SP8 confocal microscope (Carl Zeiss GmbH, Oberkochen, Germany). Quantitative analyses of acquired images were performed using ImageJ software (1.48v; National Institutes of Health, Bethesda, MD).

### Comparison to zebrafish lateral plate mesoderm (LPM)

The upregulated genes in CEP83^-/-^ cells at day 7 and at day 25 were compared with the top 20 orthologous genes identified in subclusters of zebrafish LPM identified by scRNA-seq (Prummel et al., 2022), as deposited on ArrayExpress (E-MTAB-9727)^33^.

### Statistical analysis

scRNA-seq was done on two biological replicates representing two different clones of CEP83-/- and control cells, respectively. All other experiments were performed using three biological replicates representing three independent clones of *CEP83^-/-^* and control cells at different time points. A common excel sheet for the genes present in both bulk RNA and scRNA sequencing were generated in R. The sheet includes in total 20894 genes and represents the TPM values of both groups (*WT* and *KO*) on day 0, day7, day 25 of differentiation. The maximal TPM (TPMmax) and the minimum TPM (TPMmin) were calculated for each gene across all samples. Highly variable genes (HVGs) were calculated based on the ratio of TPMmax and TPMmin. For heatmaps and PCA analysis, the top 1000 HVGs were plotted with selection of TPMmax >2 for each gene. Deregulated (upregulated and downregulated) genes between *WT* and *KO* groups were selected with expression criteria of TPM >2, fold change > 1.5, and *P*-value calculated on log10 TPM < 0.05. The unpaired 2-tailed t-test was used to compare two groups. All graphs were generated using GraphPad Prism 7.04 (GraphPad Software, San Diego, CA). Data are presented as mean ± SD.

## Results

### CEP83 is essential for the differentiation of human induced pluripotent stem cells into kidney cells

To investigate the effect of *CEP83* loss on the differentiation of hiPSCs into intermediate mesoderm (IM) kidney progenitors, we applied CRISPR-Cas9 technology to induce a null mutation in the *CEP83* gene in hiPSCs **(Figure 1A)**. Three hiPSCs clones designated *CEP83^-/-^* (*KO1*, *KO2*, and *KO3*) carried deletions within *CEP83* exon 7, each of which led to an induction of a premature stop codon resulting in a predicted truncated protein (**Figure 1B-D and Figure 1-figure supplement 1A)**. These clones exhibited a complete loss of CEP83 protein by immunoblotting (**Figure 1E**). Three wildtype clones were derived as controls (*WT1*, *WT2*, and *WT3*). All six clones were morphologically indistinguishable (by brightfield microscopy), and had similar overall gene expression profiles (by bulk RNA-seq and qRT-PCR), including pluripotency and lineage marker expression **(Figure 1-figure supplement 1B, C, and Figure 1-figure supplement 2A, B)**. In KO clones, the anticipated altered transcripts of CEP83 were detectable based on bulk RNA-seq (data not shown). Single nucleotide polymorphism (SNP) - analysis confirmed identical karyotypes of all six clones **(Figure 1-figure supplement 2C)**.

Together, these findings confirmed successful deletion of CEP83 in iPSCs without any overt direct cellular phenotypic consequences. We applied a 7 day monolayer protocol using timed application of WNT and FGF agonists as reported by Takasato et al^17^ to differentiate *WT* and *KO* hiPSCs into IM kidney progenitors^14–16^ (**Figure 2A**).

**Figure 2:**
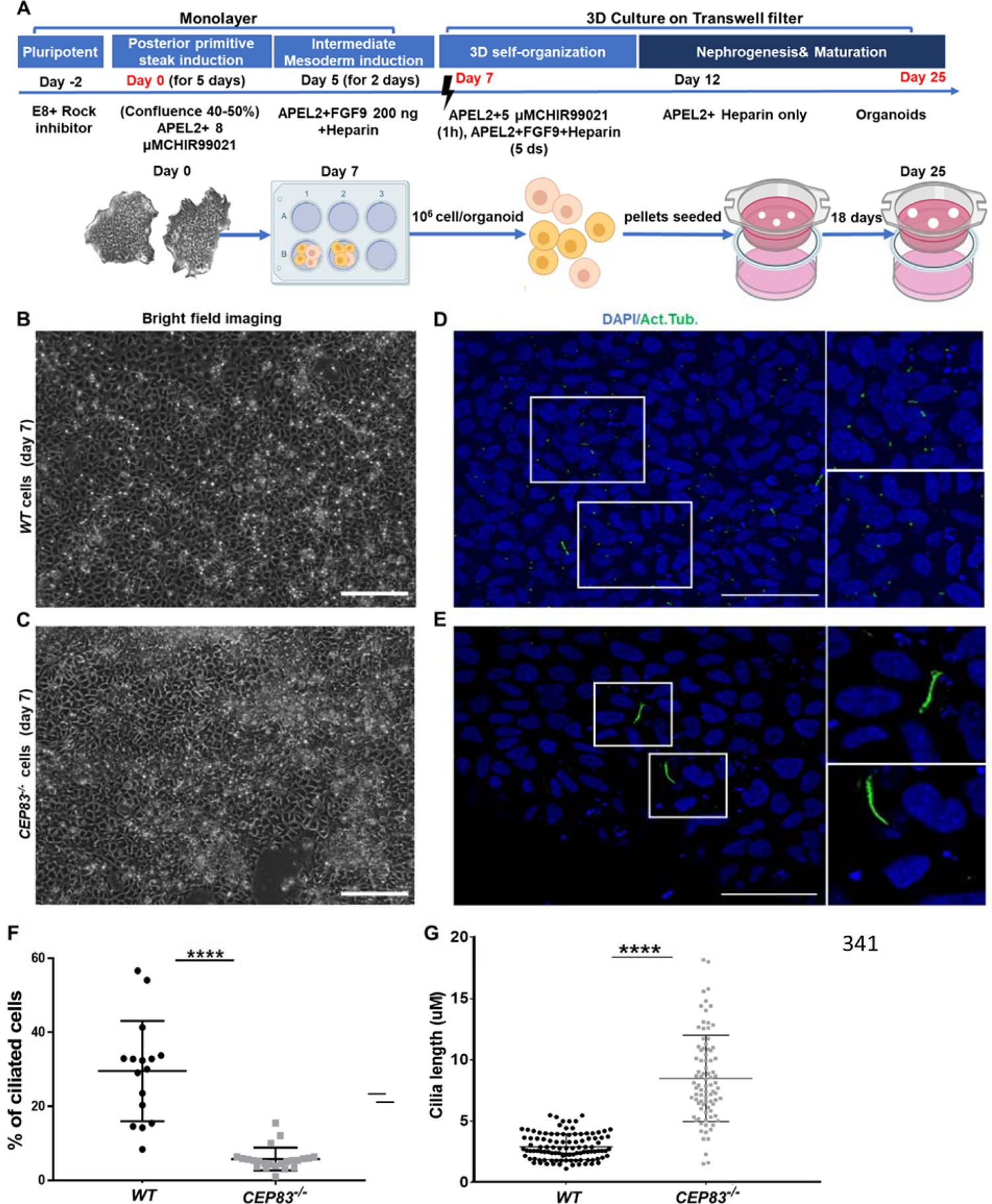
Differentiation of CEP83^-/-^ hiPSCs to intermediate mesoderm cells (day 7) is associated with defective ciliogenesis. (A) The schematic diagram illustrates the applied differentiation protocol of hiPSCs, as previously described by Takasato *et al*.^34^. (B-C) *WT* and *CEP83^-/-^* cells on D7 of differentiation did not show overt morphological differences by brighfield microscopy. (D-E) Representative images of *WT* and *CEP83^-/-^* cells on D7, immunostained for acetylated tubulin (green) and nuclei (DAPI, blue), revealing fewer and elongated cilia in *CEP83^-/-^* cells. (F) Quantitative analysis of the percentage of ciliated cells in *WT* and *CEP83^-/-^* cells (D7). (G) Quantitative analysis of the ciliary length in *WT* and *CEP83^-/-^* cells (D7). n = 3 clones per group. *\*\*\*\*P* < 0.0001. Bar = 50 μm. See **Figure 2-figure supplement 1**.

After 7 days of culture (D7), *WT* and *KO* cells exhibited an indistinguishable morphology by bright field microscopy (**Figure 2B, C)**. Immunostaining for acetylated tubulin, however, indicated abnormal primary cilia formation in *CEP83*-deficient cells (**Figure 2D, E).** The number of ciliated cells was reduced from approximately 30% (in *WT* clones) to less than 10% (in *KO* clones) (**Figure 2F**). Among ciliated cells, the length of cilia was increased from 2-5 µm (in *WT* clones) to 5-13 µm (in *KO* clones) (**Figure 2G**). This indicated that *CEP83*^-/-^ hiPSCs differentiated towards IM progenitors exhibited ciliary abnormalities. To analyze the induced IM kidney progenitor cells functionally, we collected D7 *WT* and *CEP83^-/-^* cells and placed them into an organoid culture system again applying timed WNT and FGF agonists to foster differentiation of mature kidney cell types, as previously reported^17^ (**Figure 2A**). Organoids harvested from *WT* clones after a total of 25 days of culture (D25) had formed patterned kidney epithelial-like structures, including NPHS1-positive glomerulus-like structures, Lotus tetragonolobus lectin (LTL)-positive proximal tubule-like, and E-cadherin (E-cad)-positive distal tubule-like structures (**Figure 3A, C, E)**. In contrast, *CEP83^-/-^* organoids at day 25 were composed of monomorphic cells with a mesenchyme-like appearance, which stained negative for an array of kidney cell markers (**Figure 3B, D, and F**).

**Figure 3:**
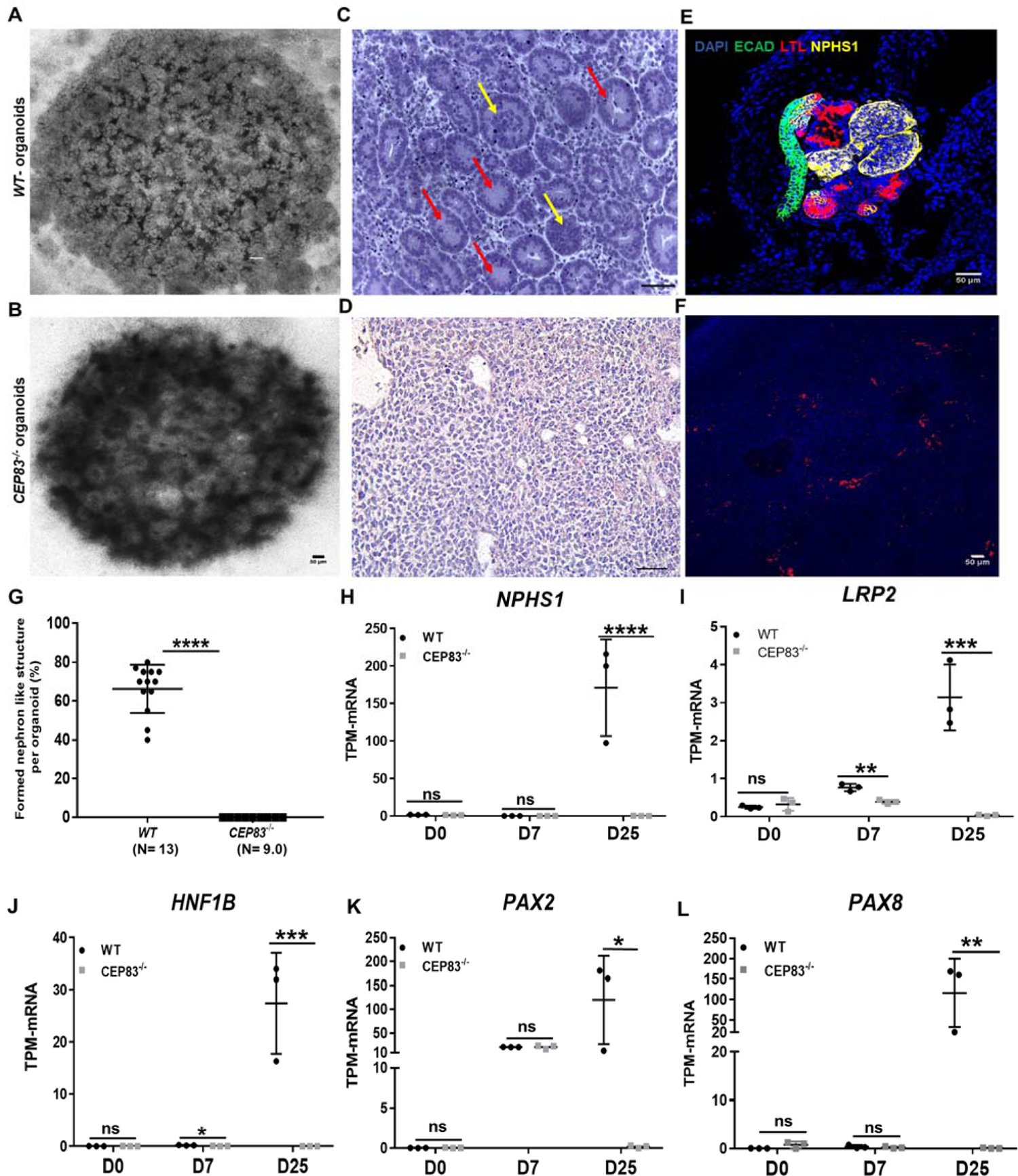
Defective kidney organoid differentiation from *CEP83*-deficient pluripotent stem cells. (A, B) Bright-field images of organoids after a total of 25 days of culture (D25) indicate formation of multiple kidney-like structures in *WT* organoids (A), whereas *CEP83^-/-^* organoids are composed of uniform clusters (B). (C, D) Representative images of hematoxylin-eosin (HE)–stained sections of organoids. *WT* organoids (C) display glomerulus-like (yellow arrows) and tubular (red arrow) components, whereas *CEP83^-/-^* organoids (D) are composed of monomorphic mesenchymal-like cells. (E-F) Whole mounting immunostaining of organoids for NPHS1 (podocyte marker), LTL (proximal tubule marker), and CDH1 (distal tubule marker) indicate segmented nephron-like structures in *WT* organoids (E) and absence of such structures in *CEP83^-/-^* organoids (F). (G) Quantitative analysis of brightfield images indicating the estimated percentage of organoid area composed of nephron like structures, organoids were collected from three different experiments. (H-L) Gene expression (transcripts per million, TPM) of *NPHP1* (H), *LRP2* (I), *HNF1B* (J), *PAX2* (K) and *PAX8* (L) in *WT* and *CEP83^-/-^* cells at the indicated time points based on bulk RNA sequencing. n= 3 clones per group. Data are mean ± SD. *\*P* < 0.05, *\*\*P* < 0.01, *\*\*\*P* < 0.001 and *\*\*\*\*P* < 0.0001. ns= not significant. Panels B, C and D: Bar = 50 μm. See Figure 3-source data 1-2. See also Figure 3-figure supplements 1-2.

Kidney epithelial-like structures formed only in *WT*, but not in *CEP83^-/-^* organoids (**Figure 3G**). Similar to the findings in day 7 cells reported above, primary cilia were found in fewer cells of *CEP83^-/-^* organoids (<5% of cells) and were abnormally elongated (**Figure 2-figure supplement 1**).

Next, bulk RNA sequencing of *WT* (*WT1*, *WT2*, *WT3*) and *CEP83^-/-^* (KO1, KO2, KO3) organoids was carried out to evaluate differential gene expression on a genome-wide level, and RT-PCR was used to validate selected genes. Hierarchical clustering of the samples indicated strong gene expression differences between *WT* and *CEP83^-/-^* samples **(Figure 3-figure supplement 1)**. Several genes associated with kidney development and kidney epithelial differentiation were differentially expressed with high expression in *WT* organoids, but showed comparatively low or absent expression in *CEP83^-/-^* organoids: these genes included kidney-specific lineage genes (*PAX2*, *PAX8*), and lineage/differentiation markers of glomerular cells (*NPHS1*, *PODXL*, *WT1*, *PTPRO*), proximal (*HNF1B*, *LRP2*, *CUBN*) and distal (*EMX2*, *MAL2*, *EPCAM*, GATA3) kidney epithelial cells. (**Figure 3H-L, Figure 3-figure supplement 1B-H, and Figure 3-figure supplement 2).** This indicated that *CEP83^-/-^* IM progenitors failed to differentiate into kidney cells, suggesting that *CEP83* function is necessary to complete essential steps in the process of differentiation from pluripotent stem cells to kidney cells.

### CEP83 deficiency associates with molecular defects of nephron progenitor cells

We next aimed to gain molecular insights into the lineage impact of *CEP83* deficiency during the course of kidney epithelial differentiation. Since no global transcriptomic differences were detectable between *WT* and *CEP83^-/-^* hiPSCs prior to differentiation (see above), we focussed on mesodermal cell stages induced at D7, which displayed mild overall gene expression differences between *WT* and *CEP83*-deficient cells as detected by bulk RNA sequencing **(Figure 4-figure supplements 1A, B, and Figure 4-figure supplement 2)**.

A marked upregulation of nephron progenitor marker genes (*GATA3*, *HOXB7*, *HOXD11*, *EYA1*)^35–42^ was observed in both *WT* and *CEP83^-/-^* cells at day 7 (**Figure 4-figure supplement 3A-D**), suggesting that the differentiation path of pluripotent *CEP83^-/-^* cells to IM nephron progenitors was largely intact. To understand the potential molecular defects at the IM stage in more detail, we performed single-cell RNA (scRNA) sequencing on D7 *WT* and *CEP83^-/-^* cells (representing two different hiPSC clones for each condition differentiated in two separate experiments). We obtained transcriptomes from 27,328 cells, representing clones *WT1* (experiment 1, 3,768 cells), *WT2* (experiment 2, 5,793 cells), *KO1* (experiment 1, 8,503 cells), and *KO2* (experiment 2, 9,264 cells). Principal component analysis (PCA) using pseudo-bulk expression data of the top 1000 highly variable genes indicated that the first major component (dimension 1, explaining 54% of expression variation) was driven by the genotype (*WT* vs. *KO*), while the second major component (dimension 2, explaining 51% of expression variation) was driven by a batch effect of the two experiments (**Figure 4A**). We combined all cells and generated a Uniform Manifold Approximation and Projection (UMAP) plot uncovering 10 different cell states/clusters (0-9; **Figure 4B**). We identified marker genes for each cluster (**Figure 4C**), indicating that clusters 1, 3 and 4 represented kidney progenitors/nascent nephrons (expressing e. g. *PAX8*, *EYA1*, *HOXB7*) in different phases of the cell cycle. Other clusters represented as-of-yet uncharacterized cell types, which was consistent with previous single cell transcriptome analyses from iPSC-derived cells induced by the same induction protocol^43, 44^. Each of the four samples (*WT1*, *WT2*, *KO1*, *and KO2*) contributed to each cluster (**Figure 4D**). We focussed on kidney progenitors (cluster 1, 3, 4) and found that a substantially lower percentage of *KO* cells (11.9-12.5%) contributed to cluster 1 when compared with *WT* cells (25.9-36.3%) (**Figure 5A**). In contrast, similar percentages of *WT* and *KO* cells were represented in kidney progenitor clusters 3 and 4 (**Figure 5B, C**). Differential gene expression analysis in these three clusters indicated significantly lower expression of kidney progenitor markers *PAX8*, *EYA1* and *HOXB7* in *KO* cells from clusters 1, 3, and 4 when compared to *WT* cells (**Figure 5D, E, F; Figure 5-figure supplement 1**). These results indicate that *CEP83* deficiency remained permissive with initial kidney progenitor induction, but that these cells exhibited mild molecular defects detectable by differential expression of kidney progenitor genes, which potentially contributed to the failure of *CEP83*-deficient cells to further differentiate towards mature kidney cell types.

**Figure 4:**
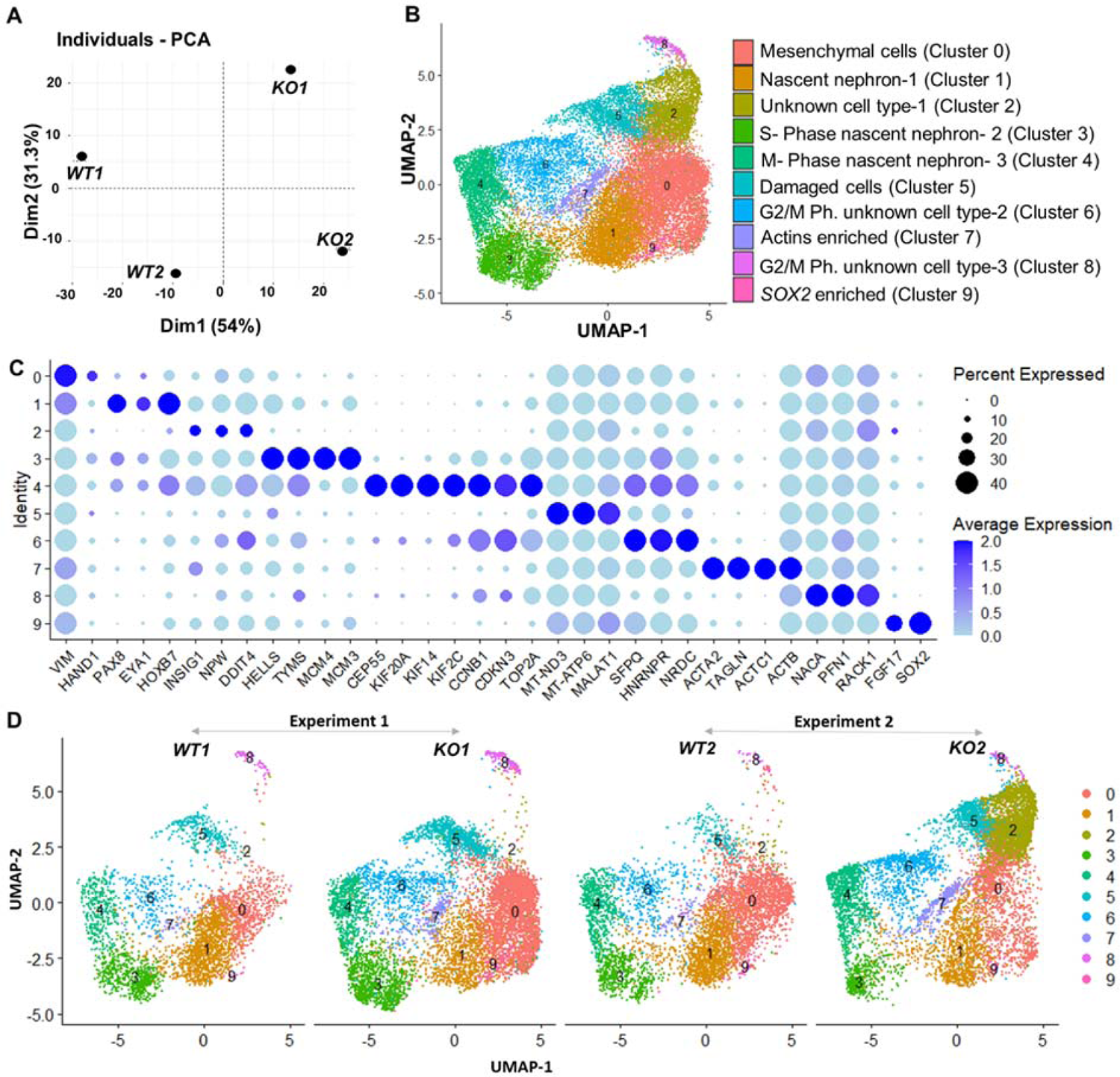
Gene expression differences of WT and CEP83^-/-^ D7 monolayers based on bulk and single cell transcriptomics. (A) Principal component analysis (PCA) of *WT* (*WT1, WT2*) and *CEP83^-/-^* (*KO1, KO2*) cells at day 7 using the average gene expression of the top highly variable 1000 genes in pseudo-bulk scRNA sequencing data. The % variation explained by each PCA axis is indicated in brackets. (B) PCA eigenvalues indicates that the principal components, Dim 1 (54%) and Dim 2 (31.3%), account for 85.3 % of the expression differences. Dim 1 separates the WT samples from the KO samples, while Dim 2 separates experiment 1 (*WT1, KO1*) from experiment 2 (*WT2, KO2*). (B) UMAP of scRNA-seq profiles from 27,328 cells from two wildtype clones (*WT1, WT2*) and two *CEP83^-/-^* clones (*KO1, KO2*) derived from two separate experiments (experiment 1: *WT1, KO1*; experiment 2: *WT2, KO2*). Unbiased clustering resulted in 10 clusters and (C) dot plot showing expression of selected marker genes of each cluster. (D) UMAP plots for *WT* and *KO* samples showing the distribution of all clusters per sample N=2 per group in B-D. See Figure 4-figure supplements 1-3. Source data is available as described in section (Data availability).

**Figure 5:**
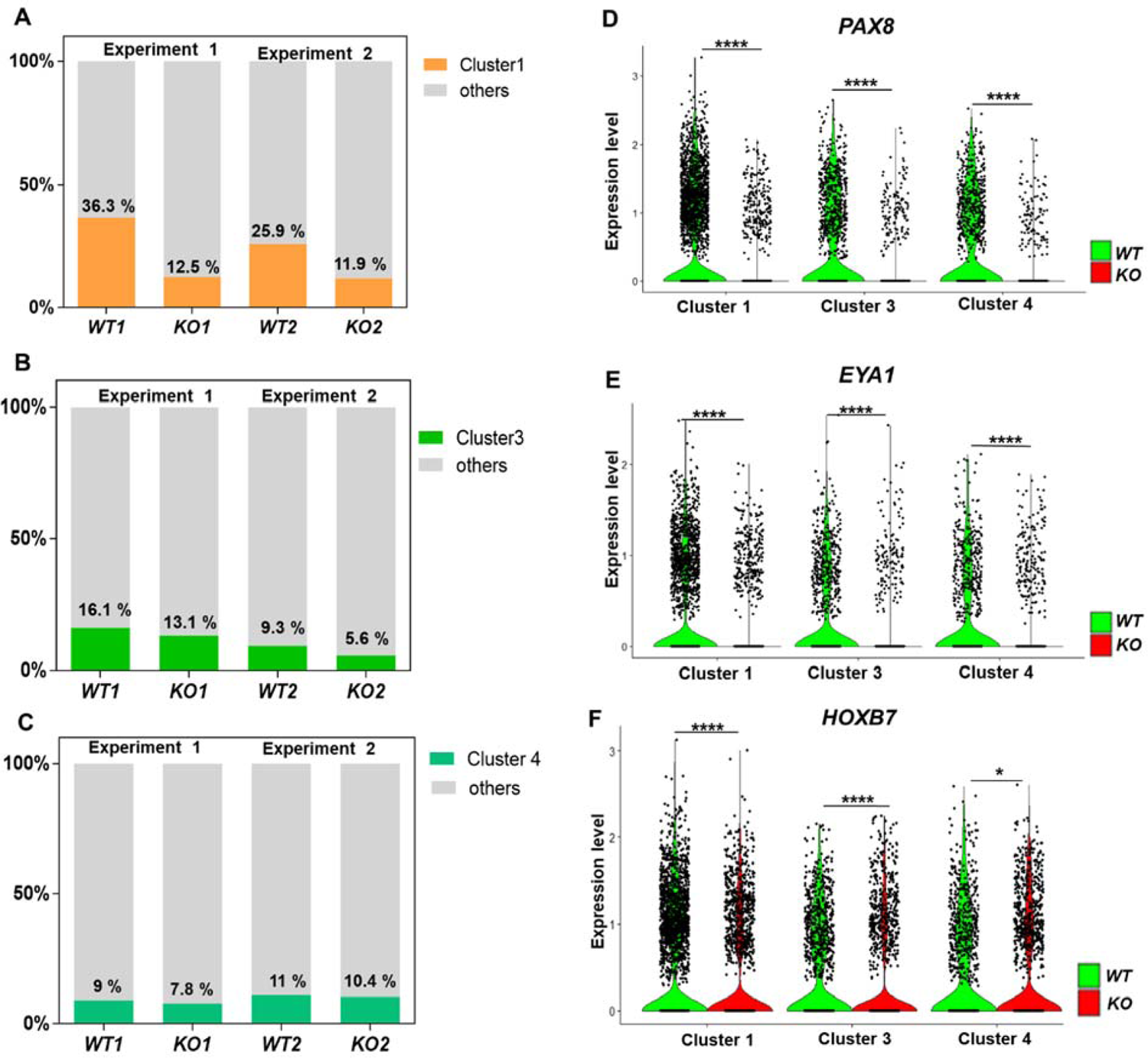
Defective kidney progenitor differentiation from *CEP83^-/-^* cells after 7 days of monolayer induction. (A, B, C) Proportions of cells from kidney progenitor clusters 1 (A), 3 (B) and 4 (C) among *wildtype* (*WT1, WT2*) and *CEP83^-/-^* (*KO1, KO2*) cells. (D, E, F) Violin plots of gene expression of kidney progenitor genes *PAX8* (D), *EYA1* (E) and *HOXB7* (F) within kidney progenitor clusters 1, 3 and 4 comparing wildtype (*WT*) and *CEP83^-/-^* (*KO*) cells. N= 2 per group. *\*P* < 0.05 and *\*\*\*\*P* < 0.0001. Figure 5-figure supplement 1. Source data is available as described in section (Data availability).

### *CEP83* deficiency promotes ectopic induction of lateral plate mesoderm-like cells followed by an expansion of cardiac and vascular progenitors

We next inspected single cell transcriptomes and bulk RNA sequencing data from D7 cells for genes that were up-regulated in *CEP83^-/-^* cells compared to WT cells. From this analysis, we observed a consistent upregulation of genes that are normally expressed in early lateral plate mesoderm (LPM), including *OSR1*, *FOXF1*, *FOXF2*, *FENDRR*, *HAND1*, *HAND2*, *CXCL12*, *GATA5*, and *GATA6*^45–69^ (**Figure 6A-I**). This suggested that *CEP83^-/-^* cells entered an aberrant differentiation path assuming a phenotype indicative of broader LPM instead of more specific IM. To further substantiate this idea, we restricted the analysis to progenitor cells of clusters 1, 3, and 4 and to cells from cluster 0, which exhibited a mesenchymal transcriptome fingerprint (see **Figure 4C**). Within each cell, we analyzed the expression of LPM markers (*FOXF1*, *HAND1*, *HAND2*, and *CXCL12*) and of more restricted IM markers (*PAX8*, *EYA1*, *and HOXB7*) (**Figure 6-figure supplement 1**).

**Figure 6:**
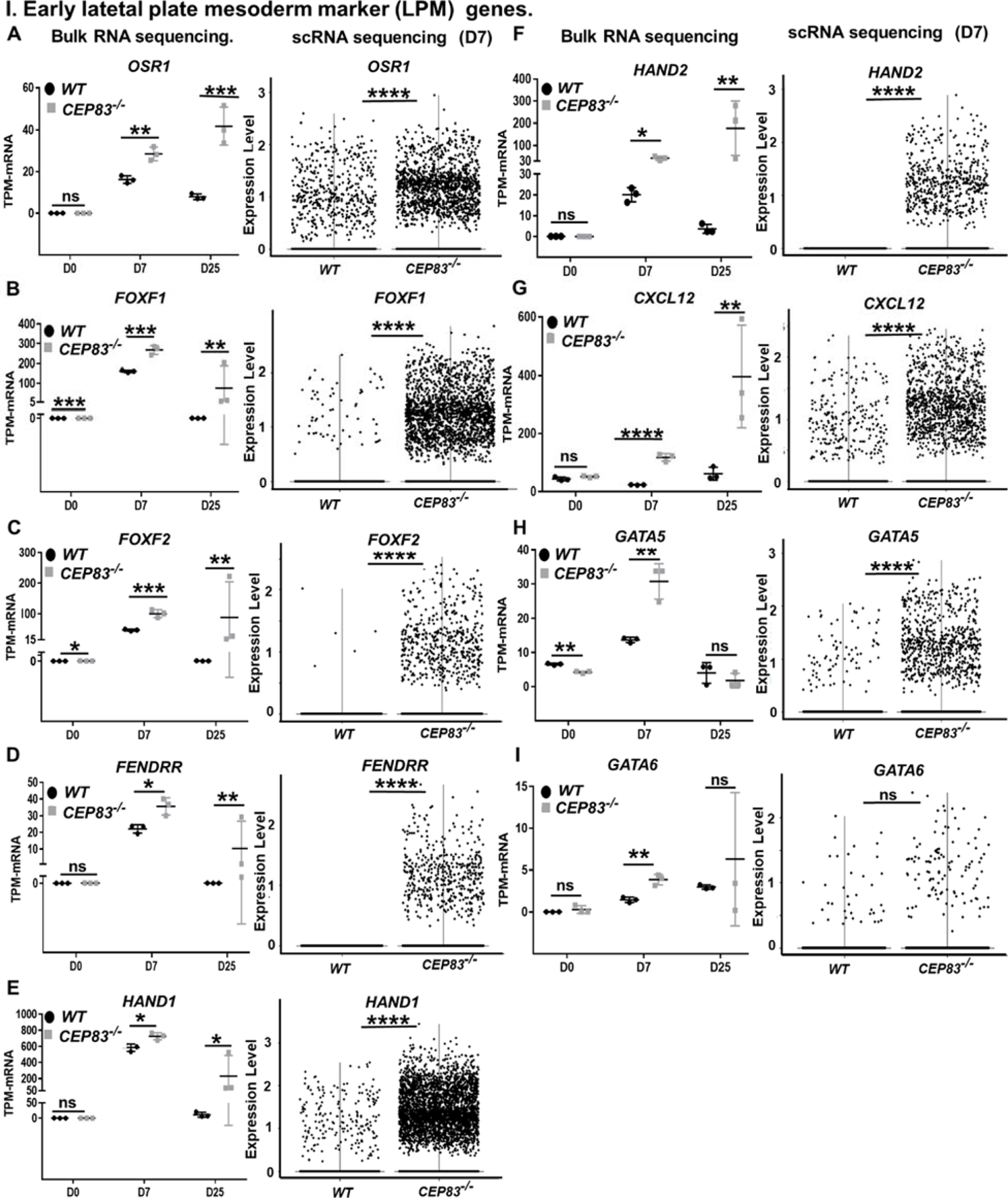
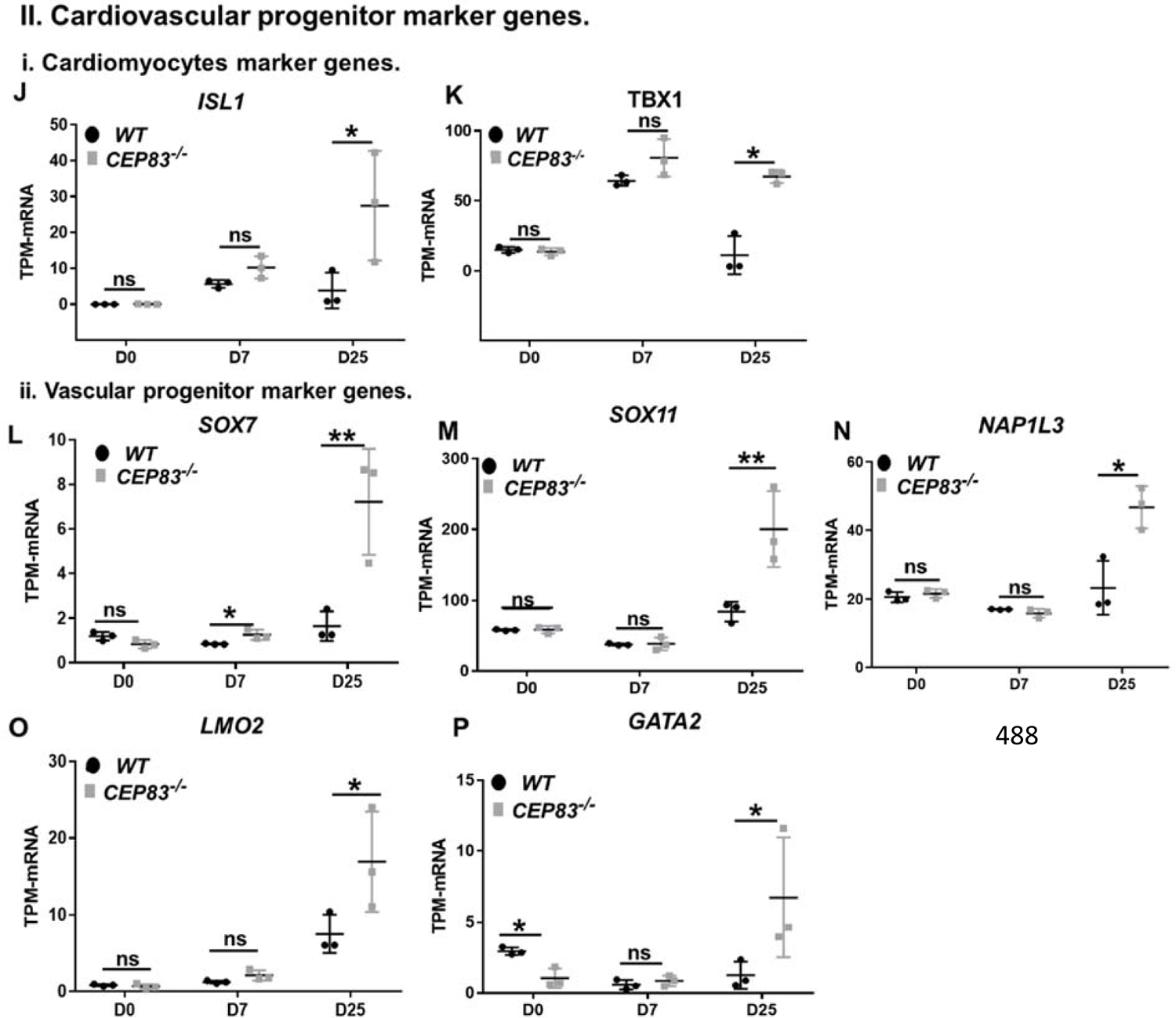
*CEP83^-/-^* cells upregulate expression of genes characteristic of early lateral plate mesoderm, cardiomyocyte progenitors and vascular progenitors. (A-I) Expression of early lateral plate mesoderm (LPM) markers *OSR1* (A), *FOXF1* (B), *FOXF2 (*C), *FENDRR* (D), *HAND1* (E), *HAND2* (F), *CXCL12* (G), *GATA5* (H), and *GATA6* (I) in wildtype (*WT*) and *CEP83^-/-^* cells at day 0 (D0), day 7 (D7) and day 25 (D25) according to bulk RNA-sequencing (left panels) and at D7 according to single cell RNA sequencing (right panels). (J-P) Expression of cardiomyocyte markers *ISL1* (J), *TBX1* (K) and vascular progenitor markers *SOX7* (L), *SOX11* (M), *NAP1L3* (N), *LMO2* (O) and *GATA2* (P) in wildtype (*WT*) and *CEP83^-/-^* cells at day 0 (D0), day 7 (D7) and day 25 (D25) according to bulk RNA-sequencing. N = 3 clones per group for bulk RNA seq. N = 2 clones per group for scRNA-seq. Expression units are mean transcripts per million (TPM) ± SD. *\*P* < 0.05, *\*\*P* < 0.01, *\*\*\*P* < 0.001 and *\*\*\*\*P* < 0.0001. ns= not significant. See Figure 6-figure supplements 1-2. See also Figure 6-source data 1. Check data availability section for other source data.

This analysis indicated that *WT* cells of these clusters exhibited an IM-like phenotype, while *KO* cells were shifted towards an LPM-like phenotype. The common IM/LPM marker *OSR1* was expressed at higher level in *KO* cells comparing to the *WT* cells.

We then inspected RNA-seq data from *WT* and *KO* organoids at day 25 for the expression of LPM genes and markers of LPM derivatives. The expression of several LPM genes (*OSR1*, *FOXF1*, *FOXF2*, *FENDRR*, *HAND1*, *HAND2* and *CXCL12*) was strongly up-regulated in *KO* cells compared to *WT* cells suggesting that an LPM-like cell pool persisted in D25 *KO* organoids (**Figure 6A-I**). To further substantiate the potential differentiation of the *CEP83*-mutant cells into broadly LPM-like cells, we compared genes that were upregulated genes in D25 organoids (in total, 397 genes) with LPM genes that were previously identified by single cell transcriptomics of sorted post-gastrulation LPM cells from developing zebrafish^33, 59, 60^. Our targeted comparison documented that *CEP83^-/-^* organoids showed significant enrichment for expression of orthologs of early LPM genes (p=0.006) (**Figure 6-figure supplement 2)**, including *OSR1*, *CXCL12*, *HAND1/2*, *KCTD12*, *PIK3R3*, and *ZBTB2*. A subset of LPM genes enriched for expression in *CEP83*-mutant cells at D25 of differentiation were indicative of cardiac or cardiopharyngeal (*ISL1*, *TBX1*) as well as of vascular progenitor (*SOX7*, *SOX11*, *NAP1L3*, *LMO2*, *GATA2*) differentiation^70–77^ (**Figure 6J-P**). Taken together, these observations document that hiPSCs without *CEP83* respond to an *in vitro* differentiation program towards kidney progenitors, yet diverge towards a broader LPM progenitor composition without significant IM instead.

## Discussion

This study indicates a novel contribution of CEP83 in regulating the differentiation path from human pluripotent stem cells to kidney progenitors. We pinpoint a stage at day 7 of intermediate mesoderm induction where *CEP83* loss of function results in a decreased nephron progenitor pool with down-regulation of critical kidney progenitor genes (*PAX8*, *EYA1*, *HOXB7*). At the same stage, genes typical of LPM specification (including *FOXF1, FOXF2, FENDRR, HAND1, HAND2*) are up-regulated (**Figure 7**). Functionally, these alterations are associated with an inability of *CEP83*-deficient cells to form kidney epithelia. Organoids derived from *CEP83*-deficient cells fail to induce any detectable nephron structures, suggesting a novel role for CEP83 during the specification of functional kidney progenitors in the mesoderm.

**Figure 7:**
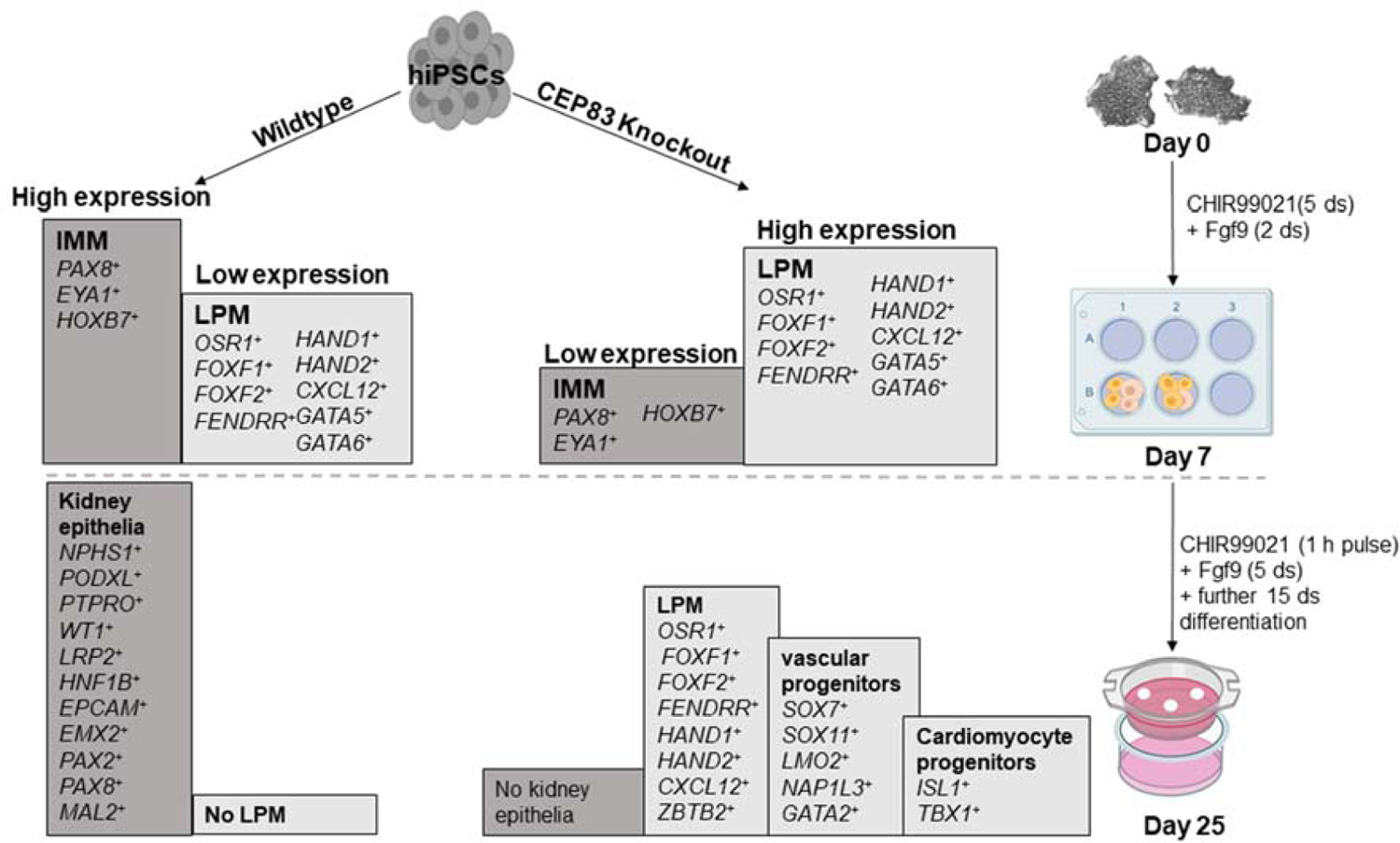
Schematic model outlining the functional differences between wildtype and CEP83 knockout cells during the course of differentiation of human pluripotent stem cells towards kidney cells. Intermediate mesoderm, IMM; lateral plate mesoderm, LPM.

Our findings are relevant to understanding cellular and molecular functions of CEP83 and might be relevant to the pathophysiology of human genetic diseases. To date, eleven patients with biallelic mutations of *CEP83* have been reported, eight of which displayed kidney phenotypes^9,78, 79^. Available kidney histologies identified microcystic tubular dilatations, tubular atrophy, thickened basement membranes and interstitial fibrosis. Extrarenal phenotypes included speech delay, intellectual disability, hydrocephalus, strabismus, retinal degeneration, retinitis pigmentosa, hepatic cytolysis, cholestasis, and portal septal fibrosis with mild thickening of arterial walls and increase in the number of the biliary canalicules on liver biopsy. Among individuals with *CEP83* mutations, all but one carried at least one missense mutation or short in-frame deletion, suggesting that *CEP83* function may have been partially preserved. One individual with presumed full loss of *CEP83* displayed a more severe phenotype with multiple organ dysfunction. It will be interesting to await future reports of additional *CEP83* mutations in humans and whether complete loss of function alleles will result in broader mesoderm defects or renal agenesis. In this regard, it is interesting that mice with a targeted homozygous loss-of-function mutation of their *CEP83* ortholog (*Cep83^tm1.1(KOMP)Vlcg^*) display midembryonic lethality (at E12.5) with evidence of severe developmental delay as early as E9.5 (https://www.mousephenotype.org/data/genes/MGI:1924298). These phenotypes are potentially consistent with a role of *CEP83* in germ layer patterning and mesoderm development, but a more detailed phenotypical characterization of *Cep83* knockout embryos would be required to substantiate this possibility.

The precise molecular and cellular mechanisms underlying our observations remain to be determined. CEP83 is a protein that is necessary for the assembly of DAPs and primary cilia formation in several cell types^4, 5, 10, 80–82^. A potential involvement of CEP83-mediated primary cilia formation in the findings reported here is suggested by obvious ciliary defects in CEP83-deficient cells at the D7 and at the organoid stage (**Figure 2D-G, Figure 2-figure supplement 1)**. These defects include reduced percentages of ciliated cells and elongated primary cilia in those cells that continue to form a primary cilium.

We observed downregulated expression of the key nephron progenitor genes *PAX8*, *EYA1,* and *HOXB7* in *CEP83^-/-^* cells at day 7, which might explain their failure to differentiate into kidney cells, since each of these genes is essential for normal kidney development ^42, 83–86^. Defects during nephron progenitor differentiation in the IM would be expected to result in severe kidney phenotypes such as renal agenesis or renal hypodysplasia. Defects of centriolar components or cilia have previously been linked to such phenotypes: in mice, centrosome amplification, i. e. the formation of excess centrosomes per cell, severely disrupts kidney development, resulting in depletion of renal progenitors and renal hypoplasia^87^. In humans, loss of KIF14, a protein necessary for proper DAP assembly and cilium formation, has been associated with kidney malformations, including renal agenesis and renal dysplasia^88–90^. Furthermore, Kif3a, a ciliary protein involved in intraflagellar transport, is necessary for normal mesoderm formation and kidney progenitor-specific defects of Kif3a have been associated with reduced nephron numbers^91, 92^. Similarly, mouse genes encoding the ciliary intraflagellar transport proteins IFT25 and IFT27 have been associated with renal agenesis or renal hypoplasia^93, 94^. Together, these studies highlight the importance of molecules involved in ciliogenesis for mesoderm and kidney progenitor development and suggest that CEP83 contributes to such processes by facilitating an early step of ciliogenesis. Nevertheless, the detailed molecular processes that link CEP83 function, cilia formation, and kidney progenitor specification remain to be determined.

The finding of various upregulated LPM markers in *CEP83^-/-^* cells starting from day 7 suggests that CEP83 function maybe involved in finetuning the balance of LPM and IM, thereby contributing to lineage decisions during mesoderm formation. Crosstalk of LPM and IM has been reported previously in zebrafish, overexpression of LPM transcription factors Scl/Tal1 and Lmo2 induces ectopic vessel and blood specification while inhibiting IM formation^95^. Furthermore, the LPM transcription factor Hand2 is critical in determining the size of the IM, while natively expressed in the IM-adjacent LPM progenitors that form mesothelia^33, 58^. Loss of Hand2 in zebrafish results in an expanded IM, whereas Hand2 overexpression reduces or abolishes the IM. Interestingly, HAND2 was among the most strongly induced transcripts in our *CEP83^-/-^* cells at day 7; connecting with the developmental role of Hand2 in IM formation, these observations suggest that HAND2 expression in CEP83-deficient cells may have contributed to the reduced numbers of nephron progenitor cells at this stage. Of note, CEP83-deficient cells at D25 expressed increased levels of LPM genes expressed in mesothelial (including *OSR1*, *CXCL12*, *HAND1/2*), cardiopharyngeal (including *ISL1*, *TBX1*), and endothelial/hematopoietic (including *TAL1*, *LMO2*, *GATA2*) progenitors^33, 60^. In sum, we propose a novel role for CEP83 in regulating the development of IM nephron progenitors, which may involve direct effects of CEP83 in the nephron progenitor differentiation program and indirect LPM-mediated effects on the IM. Future studies are warranted to delineate the molecular and cellular mechanisms underlying CEP83 function in LPM and specifically IM patterning.

## Acknowledgements

We thank Tatjana Luganskaja for excellent technical support. This work was supported by grants to K.M.S.-O. from the Deutsche Forschungsgemeinschaft (DFG; SFB 1365, GRK 2318 and FOR 2841), by stipends to F.M. by the Egyptian government, by the Urological Research Foundation (Berlin), a Swiss National Science Foundation postdoctoral fellowship to J.K.-R., and the University of Colorado School of Medicine, Anschutz Medical Campus, and the Children’s Hospital Colorado Foundation to C.M..

## Data availability

All data supporting the findings of this study are available within the article and its supplementary files. Source data files have been provided for Figures 1 to 6. Sequencing data have been deposited in GEO at https://www.ncbi.nlm.nih.gov/geo/query/acc.cgi?acc=GSE205978 (Reviewers TOKEN: mzkfymcwzzwptib).

## Supplemental Figures

**Figure 1-figure supplement 1:**
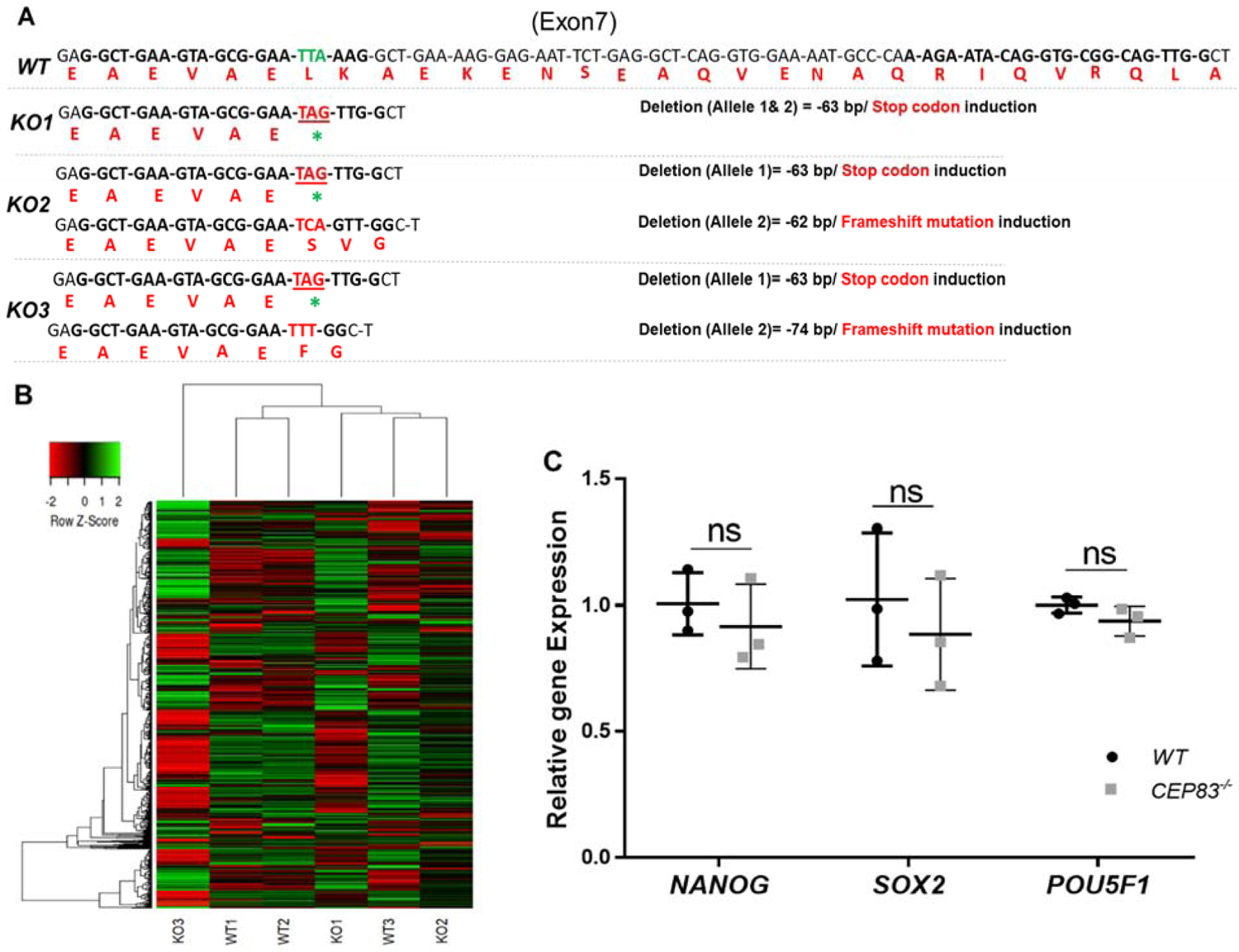
*CEP83^-/-^* hiPSCs retain global iPCS cell gene expression signatures and express pluripotency markers. (A) Alignment of the modified *KO* clones mRNA and expected amino acids sequences with *WT* revealed induction of stop codon on both strands of *KO1* clone. While *KO2* clone shows induction of stop codon on one allele and frameshift mutation within the second allele with 62 bp deletion. *KO3* clone sequence shows induction of stop codon on one allele and frameshift mutation with 74 bp deletion in the second allele. (B) Heatmap showing the expression of the top 1000 highly variable genes (see method, with a selection of TPM ≥ 10) within *WT* (*WT1, WT2*, and *WT3*) and CEP83^-/-^ hiPSCs (*KO1, KO2*, and *KO3*) clones. Unbiased hierarchical clustering of clones indicates that gene expression similarity is not driven by WT or KO status. (C) RT-PCR shows no significant differences in the expression of pluripotency markers *NANOG*, *SOX2*, and *POU5F1* between *WT* and *CEP83^-/-^* hiPSCs. TPM, Transcripts Per Million. n = 3 hiPSCs clones per group. Data are mean ± SD. ns, not significant.

**Figure 1-figure supplement 2:**
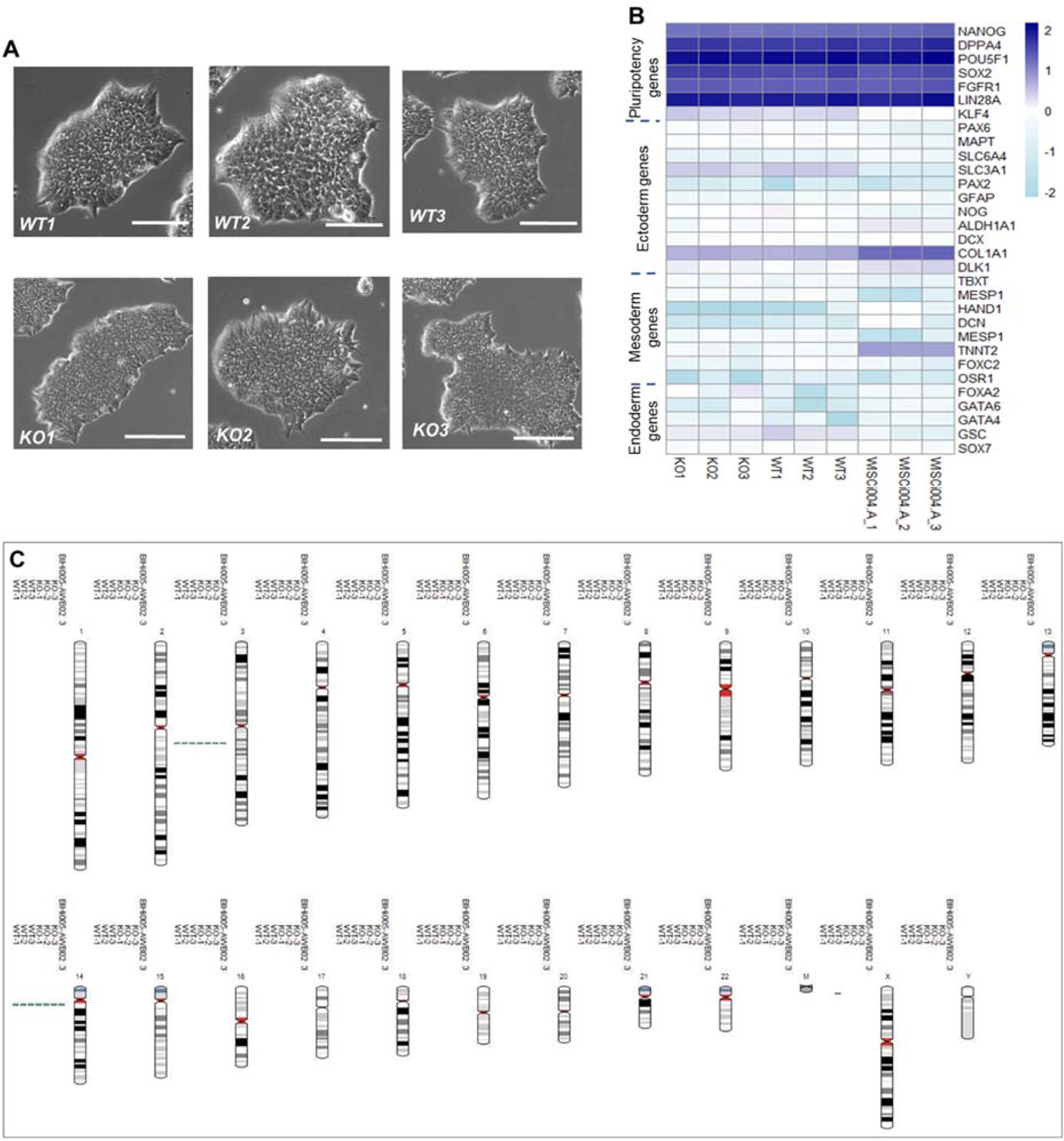
Phenotypical, molecular and genetic characterization of *CEP83^-/-^* hiPSCs versus the wildtype hiPSCs. (A) *CEP83^-/-^* hiPSCs clones (*KO1, KO2*, and *KO3*) show similar morphology to the *WT* clones (*WT1, WT2*, and *WT3*) under the bright field microscope, scale bar= 200 µm. (B) Using bulk RNA sequencing data, TPM values for marker genes for pluripotency, ectodermal, mesodermal, and endodermal cells were plotted across the samples *(KO1, KO2, KO3, WT1, WT2*, and *WT3*). In addition, gene expression of the 6 samples was compared to three wildtype hiPSCs (WISCi004-A, also referred to as IMR90-4 iPS derived from female lung fbroblasts) that were previously published^1^. (C) The three *WT* clones, three *KO* clones, and the parental population were karyotyped using single nucleotide polymorphism (SNP) - analysis, demonstrating unaffected integrity of karyotypes. Two aberrations (one gain on Chr3 and one gain on Chr14) present in BIHi005-A were previously reported (https://hpscreg.eu/cell-line/BIHi005-A, Berlin Institute of Health Stem Cell Core Facility).

**Figure 2-figure supplement 1:**
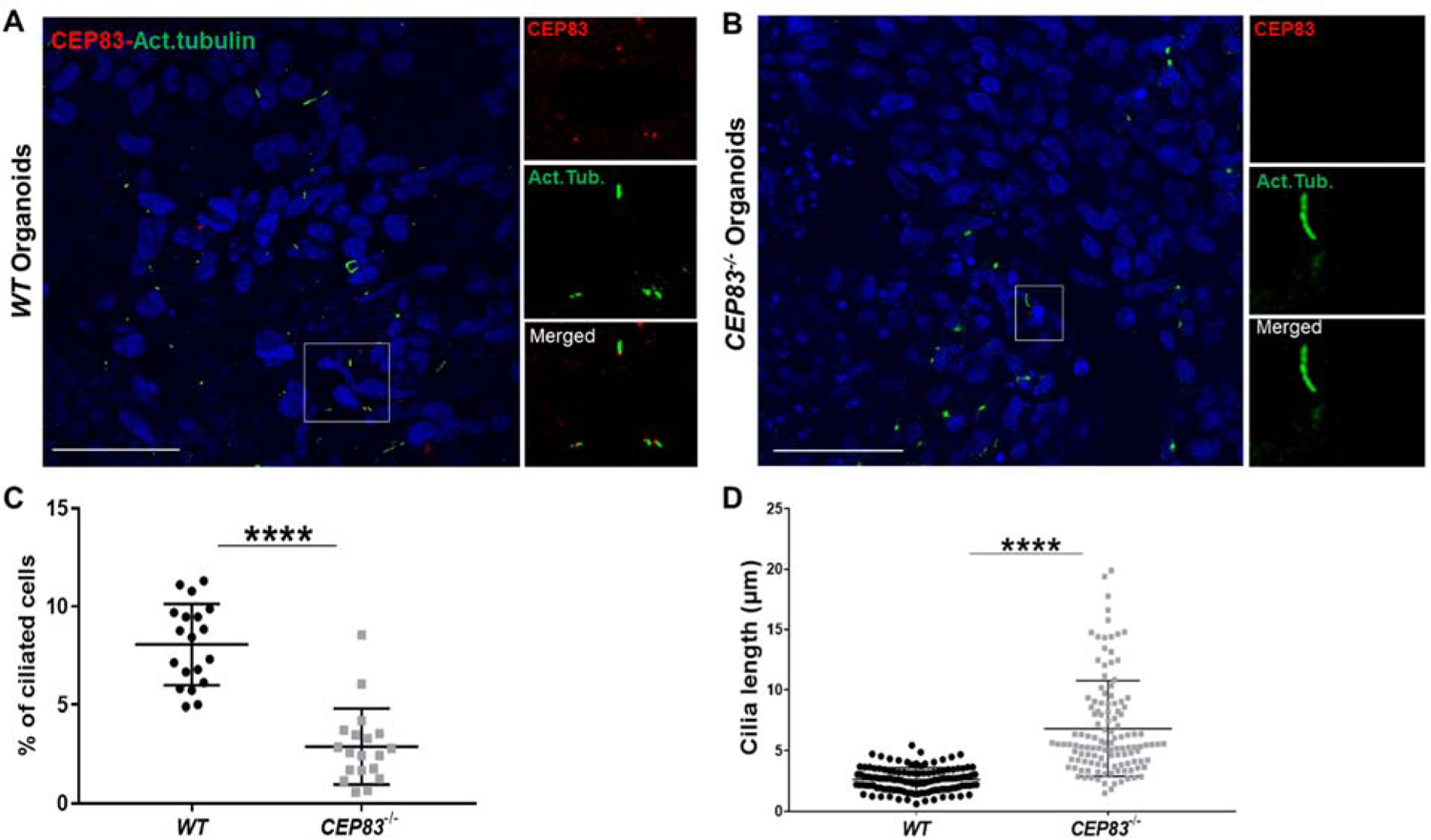
Loss of *CEP83* in organoids results in defective ciliogenesis. (A) Immunofluorescence staining of WT and CEP83^-/-^ organoids for acetylated tubulin (green), CEP83 protein (red), and nuclear staining (DAPI). Note CEP83 localization at the base of the cilium in WT organoids. (B) Quantitative analysis of ciliated cells showing downregulation of the number of ciliated cells in CEP83^-/-^ organoids, associated with longer cilium formation (C). n = 3 clones per group. Data are mean ± SD. *\*\*\*\*P* < 0.0001. Panel A: Bar = 50 μm.

**Figure 3-figure supplement 1:**
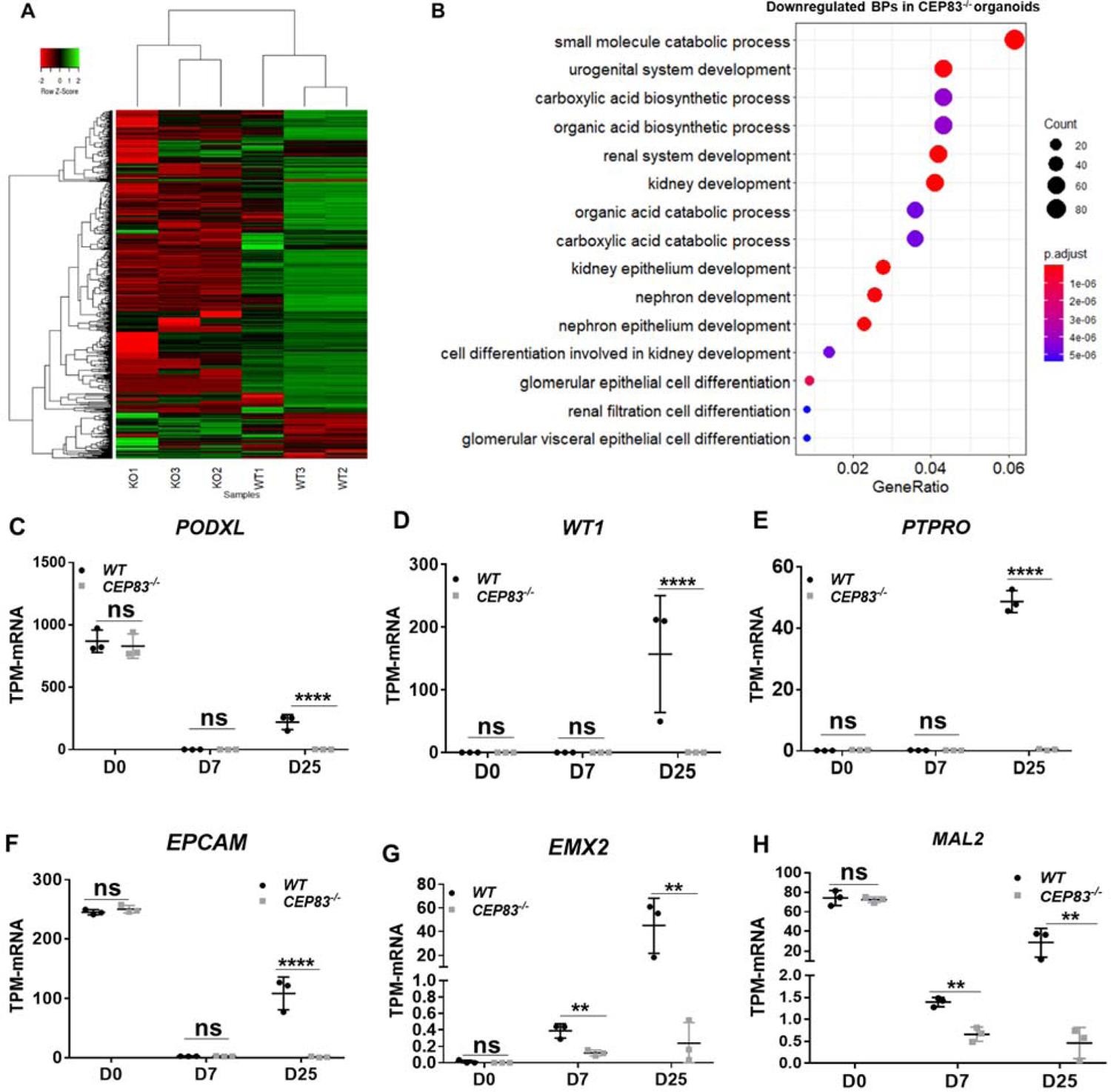
Bulk RNA-sequencing of organoids differentiated for 7+ (18) days indicates marked differences in global gene expression in *CEP83^-/-^* (*KO1-KO3*) compared to wildtype (*WT1-WT3*) organoids. (A) Heatmap displaying the expression of the top 1000 highly variable genes (see methods, TPM ≥ 10) within *WT* (*WT1, WT2, WT3*) and *CEP83^-/-^* (*KO1, KO2, KO3*) organoids. Hierarchical clustering of clones indicating that global gene expression is profoundly different in *WT* and *KO* organoids. Ontology analysis of the biological processes (BPs) using the top 100 downregulated genes (based on fold change values) in *CEP83^-/-^* organoids (TPM >2, fold change > 1.5, *P*-value calculated on log10 TPM < 0.05) using DOSE and cluster profile packages in R^2^. The analysis shows downregulation of many biological processes associated with kidney development in *CEP83* mutated organoids, as shown in the dot plot (B). Bulk RNA sequencing shows downregulation of specific renal epithelial cells marker genes at day 25, including (C-E) *PODXL*, *WT1,* and *PTPRO* for podocytes. (F-H) *EPCAM*, *EMX2*, and *MAL2* marker genes for the distal nephron precursor cells. n = 3 clones per group. Data are mean ± SD. *\*P* < 0.05, *\*\*P* < 0.01, and *\*\*\*\*P* < 0.0001. ns= not significant.

**Figure 3-figure supplement 2:**
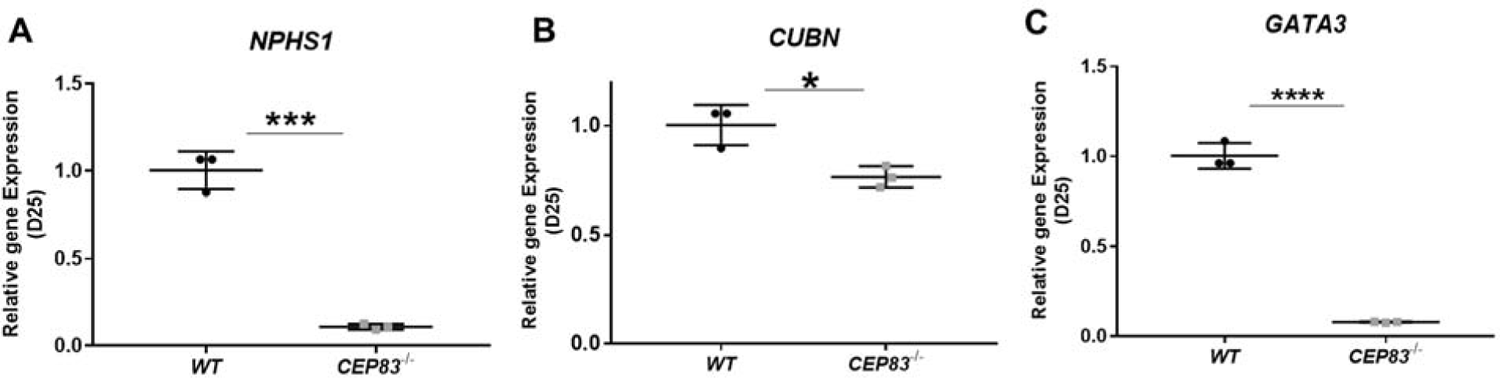
RT-PCR shows that the expression of some nephron epithelial markers, including *NPHS1* (Podocytes), *CUBN* (Proximal tubules), and *GATA3* (distal tubules and collecting duct) was significantly downregulated in *CEP83^-/-^* organoids. n = 3 clones per group. Data are mean ± SD. *\*P* < 0.05, *\*\*\*P* < 0.001 and *\*\*\*\*P* < 0.0001.

**Figure 4-figure supplement 1:**
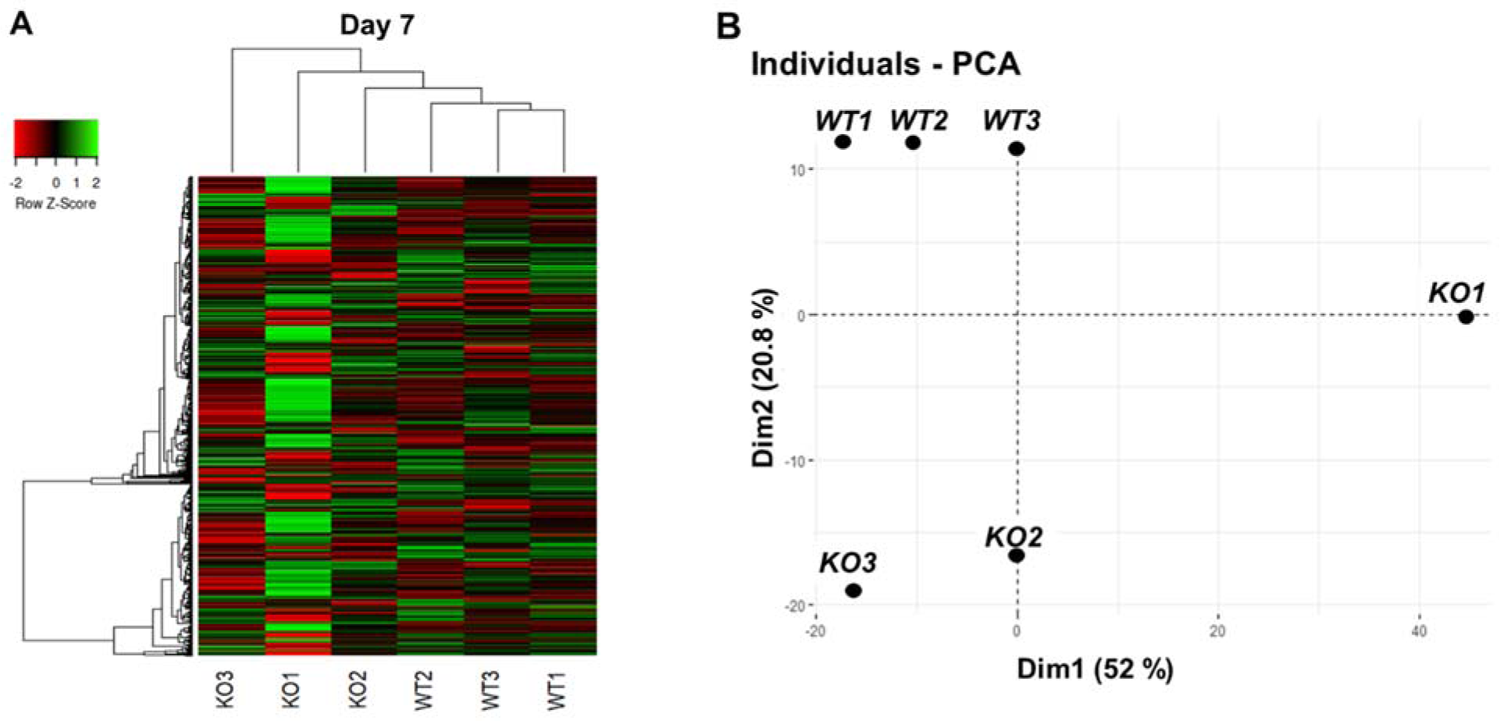
Bulk RNA sequencing shows mild overall gene expression differences between *WT* and *CEP83*-deficient cells at day 7 of differentiation. (A) Heat map of bulk RNA-seq data showing the most highly variable 1000 genes (see methods, maximum TPM ≥ 10) within wildtype (*WT1, WT2*, and *WT3*) and CEP83^-/-^ (*KO1, KO2*, and *KO3*) clones at day 7 of differentiation. Unbiased hierarchical clustering of clones separates *CEP83^-/-^* and *WT* transcriptomes. (B) Principal component analysis (PCA) of *WT* (*WT1, WT2, WT3*) and *CEP83^-/-^* (*KO1, KO2, KO3*) cells at day 7 using the average gene expression of the top highly variable 1000 genes in bulk RNA sequencing data. The % variation explained by each PCA axis is indicated in brackets. PCA eigenvalues indicate that the principal components, Dim 1 (52%) and Dim 2 (20.8%), account for 85.3 % of the expression differences. Dim 1 separates the *KO1* sample from the other samples, while Dim 2 separates experiment 1 (*WT1, WT2, WT3*) and (*KO1, KO2, KO3*)

**Figure 4-figure supplement 2:**
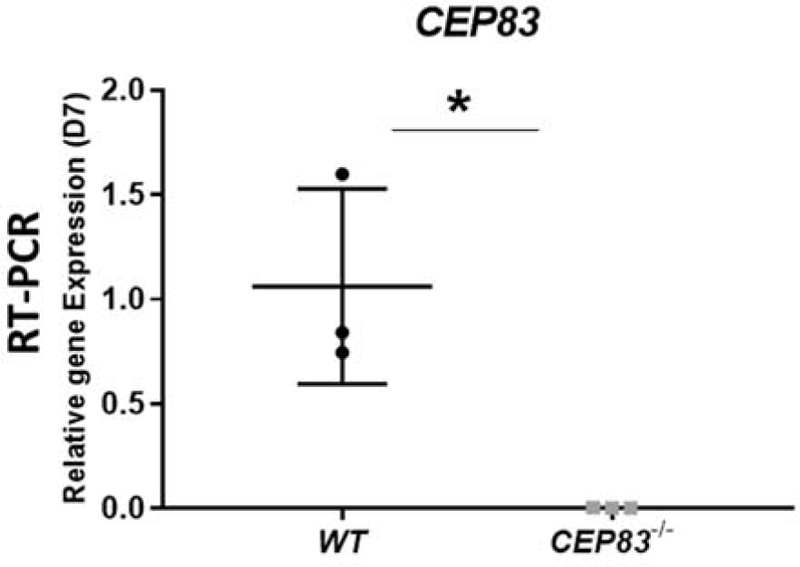
mRNA expression of *CEP83* was significantly downregulated in the *CEP83^-/-^* clones at day 7. The expression was investigated in bulk RNA seq data and confirmed by RT-PCR. n = 3 clones per group. Data are mean ± SD. *\*P* < 0.05

**Figure 4-figure supplement 3.**
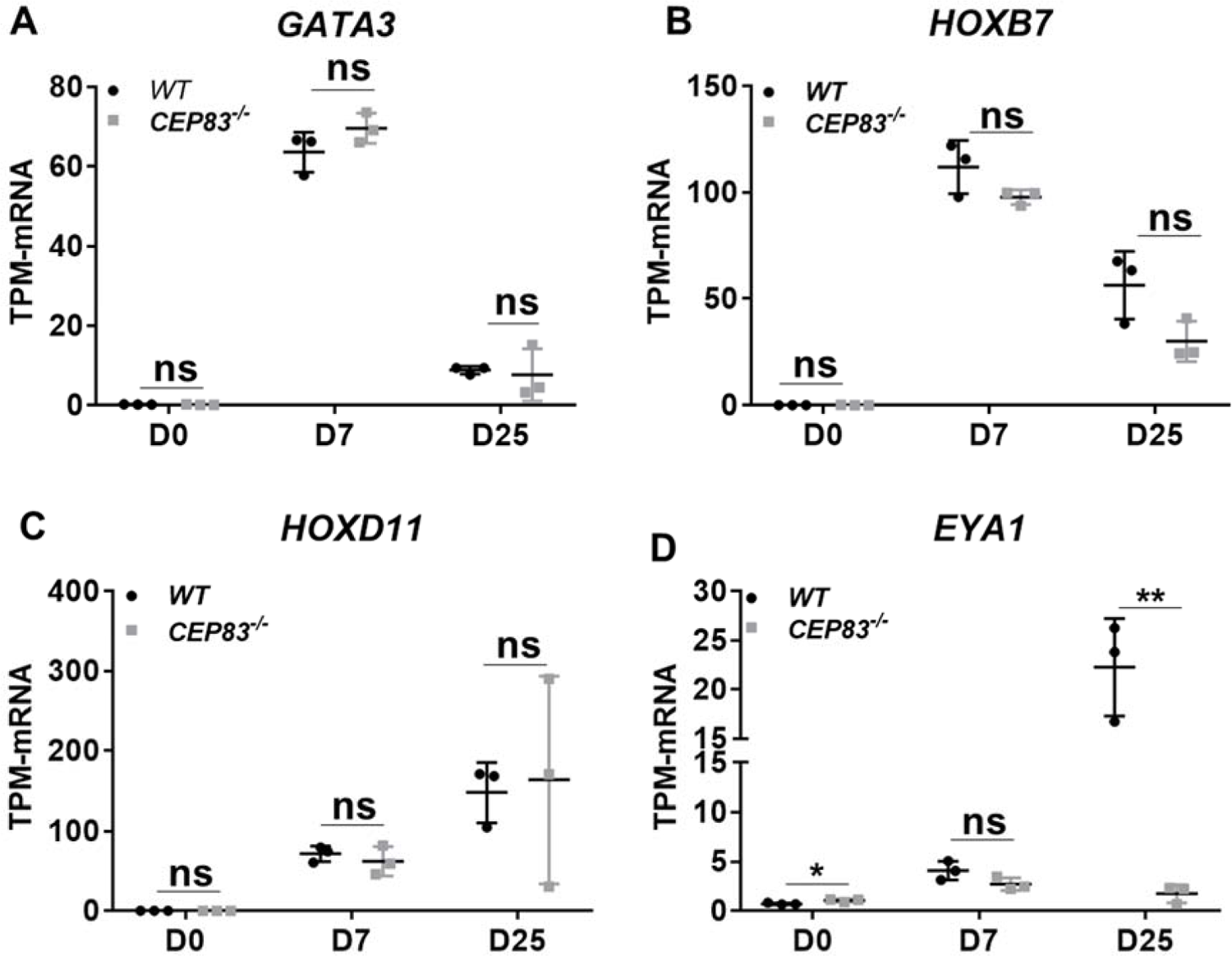
Expression of intermediate mesoderm marker genes in *WT* and *CEP83^-/-^* hiPSCs after 7 days of differentiation in a monolyer culture. (A-B) using Bulk RNA sequencing data, the expression of Ureteric bud (UB) marker genes including *GATA3*, and *HOXB7* shows no significant change between WT and mutated cells at day 0, 7, and 25. While, (C-D) MM marker genes including *HOXD11* and *EYA1* show no significant difference between *WT* and *CEP83^-/-^* cells at day 0, 7, 25 except *EYA1* show significant downregulation in the mutated cells at day 25. n = 3 clones per group. Data are mean ± SD. *\*\*P* < 0.01. ns= not significant.

**Figure 5-figure supplement 1:**
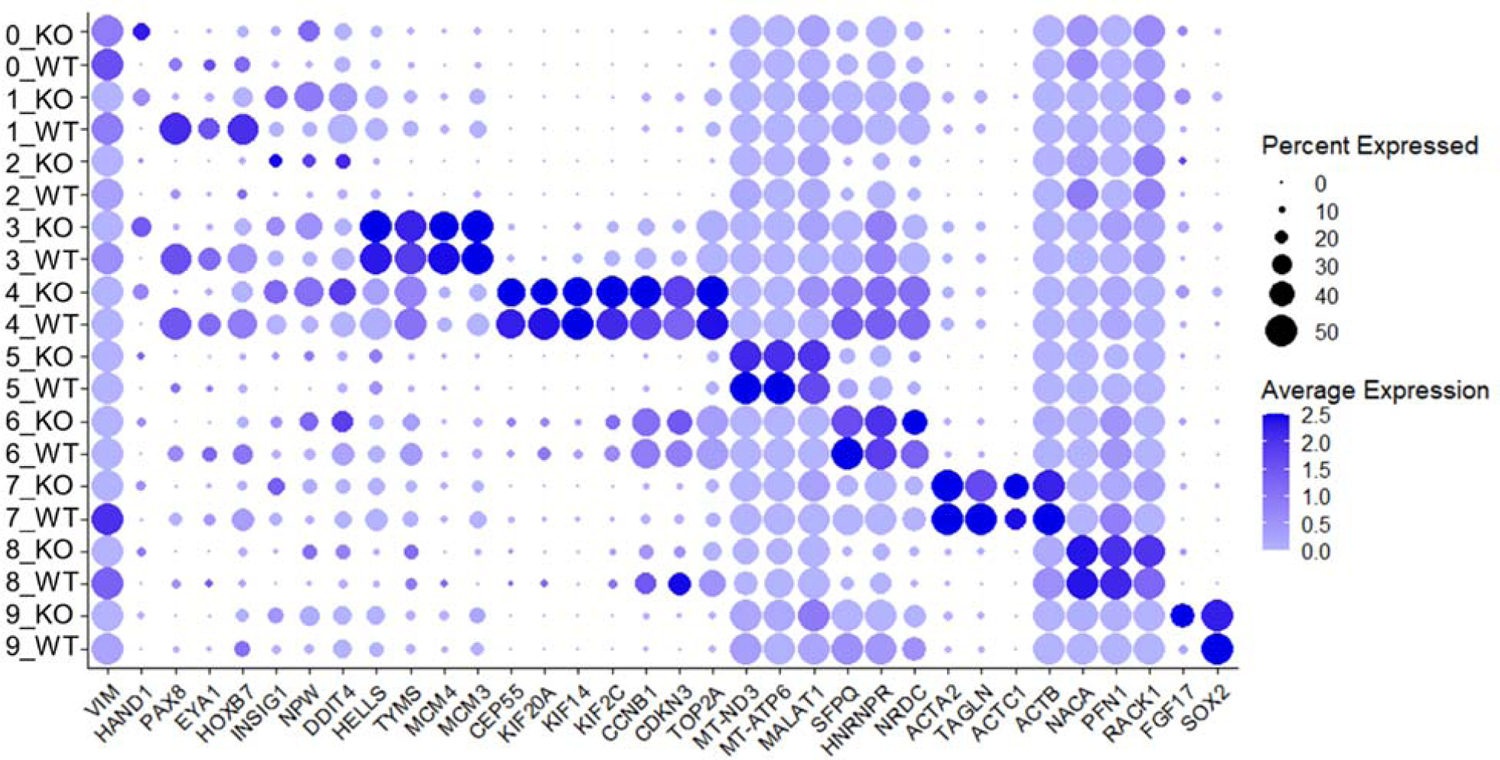
Dotplot shows the expression of marker genes of each cluster and the splitted expression per group (*WT* and *KO*). Cluster 0 (Mesenchymal cells) expresses mesenchymal genes, including *VIM* and *HAND1*. *VIM*, vimentin encodes an intermediate filament protein that plays a role in cytoskeleton organization^3,4^. The basic Helix-Loop-Helix (*bHLH*) transcription factor *HAND1* is expressed in the lateral plate mesoderm populations, in the developing heart, and in a subset of neural crest cells^5-7^. Thus, interestingly, the expression of *HAND1* is mainly represented by the *KO* cells in this cluster. Cluster 1 (Nascent nephron-1) shows upregulation of the nascent nephron marker genes, including *PAX8, EYA1*, and *HOXB7*. *PAX8* is a member of the *Pax2/5/8* family and is expressed during pro-, meso-, and metanephros development^8-10^. Interestingly, *PAX2-PAX8* double-mutant mice embryos exhibited impaired kidney development^11^. Deficiency of human *EYA1*, a homolog of the Drosophila melanogaster gene eyes absent (eya), results in an inherited disorders branchio-oto-renal (BOR) syndrome in human with or without kidney defects^12, 13^. *EYA1* knockout in mice results in complete renal agenesis^14^. *HOXB7* is one of HOX genes and is expressed in the mesonephros, ureter, and collecting system. Thus, *HOXB7* plays an essential role in kidney development. Overexpression of HOXB7 in mice causes renal duplications^15, 16^. The expression of *PAX8*, *HOXB7*, and *EYA1* was observed additionally in clusters 3 and 4. Cluster 2 (Unknown cell type-1) is expressing *INSIG1*, *NPW*, and *DDIT4*. The insulin induced gene 1 (*INSIG1*) encodes a protein that mediates feedback control of cholesterol synthesis^17^. Neuropeptide W (*NPW*) is a gene that encodes peptides that bind and activate two G-protein coupled receptors: *GPR7* and *GPR8* in the central nervous system^18^. DNA damage inducible transcript 4 (*DDIT4*) gene is expressed under stress turning off the metabolic activity triggered by the mammalian target of rapamycin (mTOR)^19^. Most of the clusters express mainly cell cycle genes of different phases. Previous studies identified the cell cycle genes for each phase^20–23^. For instance, cluster 3 (S-Phase nascent nephron-2) mainly expresses the S-phase marker genes, including *HELLS, TYMS, MCM4*, and *MCM3* in addition to the nascent nephron marker genes. Cells of cluster 4 (M-Phase nascent nephron-3) express genes of the M-phase, including *CEP55, KIF20A, KIF14*, and *KIF2C*^24^. Besides, cluster 4 expresses the nascent nephron marker genes. In contrast, cluster 5 (damaged cells) includes the damaged cells that mainly express mitochondrial genes expression as *MT-ND3, MT-ATP6*, and *MALAT1*, and show the highest mitochondrial percent compared with other clusters. While, Clusters 6 and 8 express proliferative genes that reach their peak of expression in G2/M phase genes, including *CCNB1, CDKN3,* and *TOP2A*^20–23^. Besides the G2/M phase marker genes, cluster 6 (G2/M Phase unknown cell type-2) expresses *SFPQ, HNRNPR*, and *NRDC*. Splicing Factor Proline and Glutamine Rich (SFPQ) is a ubiquitous and abundant RNA-binding protein with multiple regulatory roles in the nucleus^25^. Heterogeneous Nuclear Ribonucleoprotein R (HNRNPR) promoted cancer cell proliferation by stabilizing the expression of *CCNB1* and *CENPF* mRNA levels and promoting transcription at the proto-oncogene c-fos^26, 27^. Nardilysin Convertase (*NRDC*) gene encodes an enzyme highly expressed in human developing adult brain^28^. We couldn’t identify the type of cells of this cluster according to its gene expression profile. Cluster 7 (Actins enriched) expresses genes that encode actin proteins, including *ACTA2, TAGLN*, *ACTC1,* and *ACTB,* which are expressed in the cytoskeleton of the cell^29, 30^. While, cluster 8 (G2/M Phase unknown cell type-3) upregulates *NACA, PFN1*, and *RACK1*. The nascent-polypeptide-associated complex alpha polypeptide (NACA) gene encodes a protein associated with basic transcription factor 3 (*BTF3*) and acts as a potent Suppressor of protein aggregation and aging-related proteinopathies^31^. Profilin 1 (PFN1) plays a crucial role in promoting actin polymerization in cells^32^. Invitro study showed that receptor inhibition for activated C kinase 1 (RACK1) could suppress cell proliferation and induce apoptosis^33^. While cluster 9 (*SOX2* enriched) is expressing *FGF17* and *SOX2*. Fibroblast growth factor 17 (*FGF17*), a gene expressed in the developing brain and involved in cerebellar vermis development^34^. Sex determining region Y-box 2 (*Sox2*) gene plays a critical role in maintaining stem cell pluripotency and differentiation of pluripotent stem cells into neural progenitor stem cells^35^. n = 2 clones per group.

**Figure 6-figure supplement 1:**
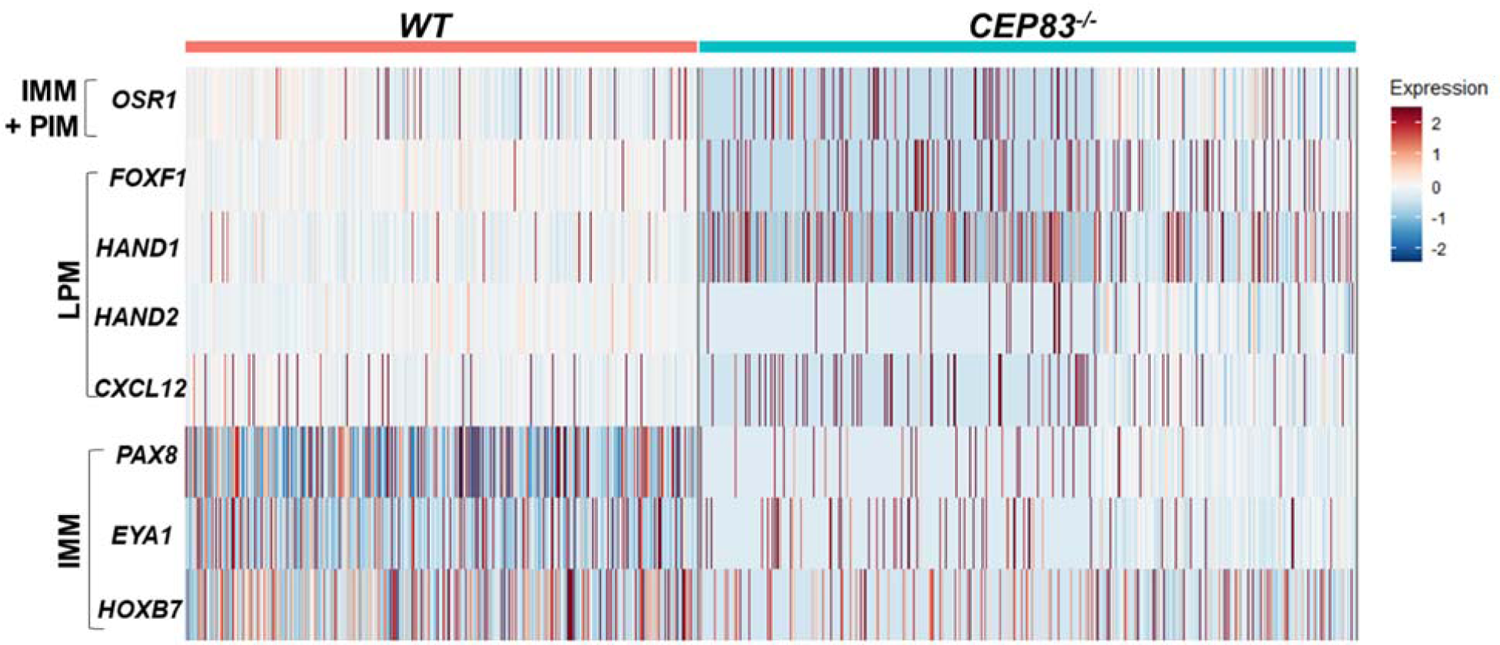
*CEP83*^-/-^ cells upregulate LPM genes in mesenchymal cells cluster and nascent nephron progenitor clusters. The heatmap shows that the log fold change of the average expression (default setting in Seurat package) of LPM and IMM genes in the WT cells (8,123 cell) and the knockout cells (10,431 cell) derived from the mesenchyme cells cluster (cluster 0) and the three nascent nephron clusters (1, 3, and 4). Scoring for cells expression for both LPM including: *FOXF1, HAND1, HAND2*, and *CXCL12,* and IMM genes including: *PAX8, EYA1*, and *HOXB7* were done in R. Each cell was scored 0-4 for LPM genes expression. 0, 1, 2, 3 and 4 mean that cell express no LPM genes, 1 gene, 2 genes, 3 genes, and 4 genes, respectively. Statistical analysis comparing between LPM scores for *WT* and *KO* cells using wilcoxon rank sum test, showed that *KO* cells significantly upregulate the expression of LPM genes. The same scoring analysis was done for the expression of the IMM genes, where the cells got score from 0 to 3 for IMM genes expression. Interestingly, the *KO* cells showed significant downregulation of the IMM genes expression.

**Figure 6-figure supplement 2:**
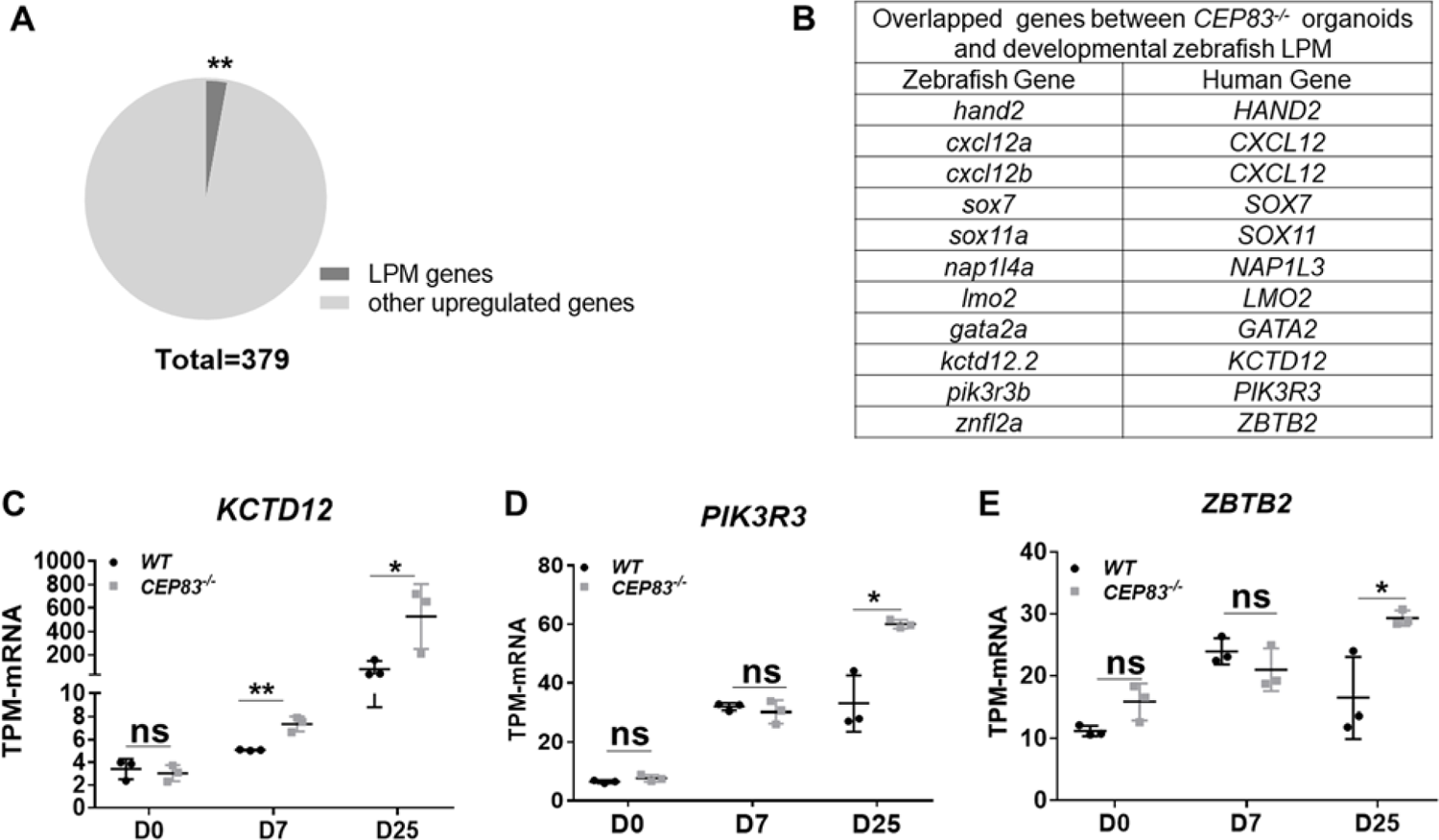
CEP83^-/-^ organoids shows significant enrichment compared to developmental zebrafish LPM scRNA data. The expression of the upregulated genes by CEP83^-/-^ organoids at day 25 were compared with the top 20 genes per cluster of zebrafish LPM scRNA data (Prummel et al., 2020). The analysis showed significant enrichment (A), with 11 zebrafish genes overlapped with 10 human genes (B). C, D, and E show the significant upregulation of three overlapped genes with zebrafish LPM including KCTD12, PIK3R3 and ZBTB2 respectively. n = 3 clones per group. Data are mean ± SD. *\*P* < 0.05, and *\*\*P* < 0.01. ns= not significant.

## Supplemental Methods

### hiPSCs Culture

We used the human iPSC cell line BIHi005-A, which was generated from a healthy donor by the Berlin Institute of Health (BIH) and supplied by the stem cell core facility at Max Delbrück Center for Molecular Medicine (Berlin). The hiPSCs were maintained in 6-well plates (Corning®, 353046) coated with Matrigel (Corning®, 354277) and cultured in Essential 8 medium (E8, Gibco-Thermo Fisher Scientific, A1517001) supplemented with 10µM Y-27632 (Rocki, Wako, 253-00513). The cells were split twice per week using EDTA/PBS or Accumax™ (Stem cell technology, 07921).

### CRISPR CAS9 Technology to generate CEP83^-/-^ hiPSCs clones

Clustered Regularly Interspaced Short Palindromic Repeats (CRISPR)-Cas9 technology was used to generate CEP83^-/-^ hiPSCs clones. Two 20 bp-long CRISPR RNAs (crRNAs) were designed using CRISPOR software^36^ to selectively target exon7: (5’-GGCTGAAGTAGCGGAATTAA-AGG-3’); (5’-AAGAATACAGGTGCGGCAGT-TGG-3’). The crRNAs were ordered from Integrated DNA Technologies (IDT). The crRNAs (IDT) were annealed in equimolar concentrations with trans-activating CRISPR RNA (tracrRNA) to form two guide RNAs (gRNA1 and gRNA2), which were then conjugated separately with Alt-R® S.p. Cas9 Nuclease V3 (1 μM concentration, IDT, 1081058) at room temperature for 1 hour to form ribonucleoprotein (RNP) complexes (RNP1 and RNP2).

One day prior to transfection, hiPSCs were split using Accumax™ solution and cultured in an equal proportion of E8 medium and StemFlex™ medium (Thermo Fisher Scientific, A3349401). The hiPSCs were transfected using a Neon transfection system (Thermo Fisher Scientific, MPK5000). Immediately before the transfection, the cells were dissociated, collected, and resuspended in Resuspension Buffer (Buffer R), that included in Neon™ transfection 100 μl kit (Thermo Fisher Scientific, MPK10025)^37^. Cells were transfected in 3 ml Electrolytic buffer (Buffer E), that included in the neon transfection kit, and by using the Neon transfection system 10µl tip. The used Neon Transfection parameters were voltage (1200 V), width (30 ms), and pulse (1). The transfected cells were cultured in StemFlex™ Medium with Rocki for 48 hours.

Then, quick DNA extraction and PCR were then done to test transfection efficiency according to the manufacturer’s instructions using Phire™ Tissue Direct PCR Master Mix (Thermo Scientific, F170S). The size of the PCR products were visualized on 1.5% agarose gel. After confirming the transfection’s success in the knockout cells, as shown in Figure 1B, the cells were dissociated and seeded at low densities for 24 hours. Then, twenty-four single cells were picked under a picking hood S1 (stem cell core facility-MDC, Buch) and cultured in StemFlex™ medium for two weeks. The clones were then tested for CEP83 mutation on the DNA level by PCR using Phire™ Tissue Direct PCR Master Mix. Finally, the selected clones were expanded and frozen in the Bambanker medium (Nippon Genetics, BB01) for further characterization. The selected clones were characterized for the mutation induction on the DNA, protein, and RNA level.

### Differentiation Protocol

We used the protocol of ***Takasato*** to differentiate the hiPSCs into nephron organoids^38^, the experiment was performed using three replicates per each group wildtype hiPSCs (*WT1*, *WT2*, and *WT3*) and *CEP83^-/-^* hiPSCs (*KO1*, *KO2*, and *KO3*). Two days prior to the differentiation, cultured hiPSCs on matrigel with 70-80% density were prepared for the differentiation. The cells wrere washed twice with 1x Dulbecco’s PBS (Thermo Fisher Scientific,14190-250), then cells were trypsinized using 1x Trypl E Select (Thermo Fisher Scientific, 12563011). Cells were incubated at 37 °C for 3 minutes. Then, DMEM/F-12 medium (Thermo Fisher Scientific, 11320-033) were added on the cells to neutralize Trypl E. The cell suspension was were mixed by pipetting (pipetting is maximum twice), then centrifuged at 300g for 5 minutes. The cell pellet was washed and resuspended in 1 ml of E8 medium. Then the cells was counted using Countess® chamber slide and Countess II Automated Cell Counter (Thermo Fisher Scientific). Then cells were centrifuged and resuspended in E8 media supplemented with 10µM Rocki. Lastly, cells were cultured on a prepared coated matrigel six well culture plates to obtain a density of 15 × 10^3^ cells per cm^2.^, and incubated overnight at 37°C CO_2_ incubator for 48 h with daily medium change.

Immediately before the differentiation, the cells were checked under microscope. Cells with 40–50% confluency were used for the differentiation. The E8 medium was changed into APEL2 medium (Stem Cell Technologies, 05270) with 5% Protein Free Hybridoma Medium II (PFHMII, GIBCO, 12040077) and 8 μM CHIR99021 (2 ml medium per a well of 6-well plate). Cells were incubated in a 37 °C CO2 incubator for **5d**, with medium refreshing every 2d. Following the CHIR99021 phase, the medium was changed into double volume of APEL2 medium (4 ml medium per a well of 6-well plate) supplemented with 200 ng/ml FGF9 (R&D, 273-F9-025) and 1 μg/ml heparin (Sigma Aldrich, H4784-250MG), and were incubated in a 37°C CO2 incubator.

On day 7 of differentiation, the cells were washed, trypsinizied with trypsin EDTA (0.05%), and incubated at 37°C for 3 min. Then, the cell suspension was transferred to a 50-ml tube containing 9 ml of MEF conditioned medium ( R&D, AR005) to neutralize the trypsin. The cells were centrifuged and resuspend in APEL2 medium. Using a hemocytometer, the cells were counted and the cell suspension was divided to achieve 1×10^6^ cells (organoid) per 1.5 ml tube. All the tubes were centrifuged at 400xg for 3 min at RT. During centrifugation, Six-well Transwell cell culture plates (Corning-Sigma Aldrich, CLS3450-24EA) were prepared by adding 1.2 ml of APEL2 supplmented with 5 μM CHIR99021 to each well. Cell pellets were picked up by using a P1,000 or P200 wide-bore tip. Pellets were carefully seeded onto the six-well Transwell membrane with minimal APEL2 medium carryover, and incubated at 37°C for 1h. Then the medium were changed into APEL2 medium supplemented with 200 ng/ml FGF9 plus 1 μg/ml heparin for further 5 days, with medium refreshing every 2 days. Finally, the medium to APEL2 medium with only heparin for further 13 days, with medium refreshing every 2 days.

### DNA isolation and Polymerase Chain Reaction (PCR)

Cultured *WT* and *CEP83^-/-^* hiPSCs were washed, scrapped gently using a cell scraper (VWR, part of Avantor, 734-2602), collected with a maximum 5×10^6^ cell/ml for proper DNA extraction. The DNA was extracted using DNeasy Blood & Tissue Kits (Qiagen, 69504) following the manufacturer’s instructions. The concentrations and quality of the DNA were evaluated using Nanodrop (Thermo Scientific, Waltham, MA; USA). To detect CEP83 expression, 200 µg DNA was amplified by a standard Polymerase Chain Reaction (PCR) using Phusion high-fidelity DNA polymerase (Biolabs, New England, M0530). The master mix was calculated according to the manufacturer’s instructions. Primers are designed using Primer3 webtool, Table S1. PCR was carried out in a thermocycler as follow: initial denaturation at 98°C for 30 sec, 35 to 40 cycles of 30 sec at 98°C, 30 sec at 63.5 °C and 30 sec at 72°C; final elongation step at 72°C for 10 min. The PCR results were checked on 1.5% agarose gel and analyzed using a BioDoc Analyze dark hood and software system (Biometra).

### RNA isolation, RNA Sequencing, and Quantitative PCR (qPCR)

Total RNA was isolated from the cells at three time points: day 0 (hiPSCs), day 7 (IMM), and day 25 (organoids) with a maximum of 1×10^7^ cells using RNAasy Mini Kit (QIAGEN, Hilden, Germany, 74104,) following the manufacture instructions. The RNA was treated with RNase-free DNase I (QIAGEN, 79254) for 15 minutes at room temperature during the extraction. The concentration, quality, and integrity of the extracted RNA were evaluated using Nanodrop (Thermo Scientific, Waltham, MA; USA), an Agilent 2100 Bioanalyzer, and the Agilent RNA 6000 Nano kit (5067-1511, Agilent Technologies). More than 0.4 μg total RNA with high integrity (more than 6.8) and high purity (OD260/280 = 1.8-2.2 and OD260/230 ≥ 1.8) were collected and sent for Illumina NovaSeq 6000RNA sequencing by Novogene. RNA-Seq library preparation and next-generation sequencing: cDNA libraries with paired-end 150 bp enriched were prepared by Novogene. Firstly the mRNA was randomly fragmentated and supplemented with oligo (dT) beads. Then cDNA synthesis were done using the random hexamers and reverse transcriptase. Secondly, second-strand synthesis was done using: a custom second-strand synthesis buffer from Illumina, deoxyribose nucleoside triphosphates (dNTPs), RNase H and E.coli polymerase I. The final obtained cDNA library was purified, terminally repaired, A-tailed, ligated to sequencing adapters, size-selected, and PCR-enriched. Quantification of library concentration was performed using a Qubit 2.0 fluorometer. Library size was measured by Agilent 2100 bioanalyzer and was quantified by qPCR (library activity > 2 nM). Libraries were sequenced on Illumina NovaSeq 6000 S4 flow cells (paired end, 150bp).

Raw data were transformed to sequenced reads, and recorded in a FASTQ file. FASTQ files were aligned to build 19 of the human genome provided by the Genome Reference Consortium (GRCh19) performed by Christian Hinze using TOPHAT2 aligner tool^39^. Up to 4 mismatches with the reference genome were accepted. Raw counts were obtained using featureCounts^40^. Mutation visualization in the knockout samples was performed using the Integrative Genomic Viewer (IGV) tool^41^. For gene expression analysis reads were normalized to the sequence length and transcripts per million (TPM) values were calculated^42^. TPM values of the samples were used to plot heatmaps and for PCA analysis based on Pearson correlation, using R (R Development Core Team, 4.0.4) 500 ng of RNA was reverse transcribed using the RevertAid First Strand cDNA synthesis kit (Thermo Scientific, K1622) according to the manufacturer’s instructions. The qPCR was carried out using the Fast Universal SYBR Green Master Mix (ROX, Roche Diagnostics, 04 913 850 001,) according to the manufacturer’s instructions. For expression analysis, relative mRNA expression levels were normalized for GAPDH mRNA expression and calculated according to the ΔΔCt method. All primer pairs were designed using the free-online primer design tool Primer3, purchased at BioTeZ (Berlin, Germany), and sequences are shown in (Table S1). Statistical significance of differences between two groups (WT and KO) was analyzed using two-sided Student’s t-test.

### Single cell RNA (scRNA) experiment. Cells isolation and preparation

The differentiated hiPSCs to intermediate mesodermal cells were collected at day 7 of the differentiation from two different experiments. The cells of the first experiment were derived from *WT1* and *KO1* differentiated hiPSCs, while the second experiment comprises the differentiated cells of *WT2* and *KO2* cells. The cells were washed twice with 1X DPBS and dissociated with Accumax for 7 min at 37 °C. Cells were centrifuged at 350 × g for 5 min, and resuspended in 1X DPBS. Then, cells were filtered a 40 µm filter (Corning, 352340), counted (10,000 cells per sample), and checked for viability using Trypan blue staining.

### Protein extraction and Immunoblotting

Protein extraction: Up to 1*10^6^ hiPSCs per sample (*WT1*, *WT2*, *WT3*, *KO1*, *KO2*, and *KO3*) were washed with cold 1xPBS, then centrifuged at 3500 g for 5 minutes. Next, the cell pellet was resuspended in pre-ice cold 100 µl of radioimmunoprecipitation assay (RIPA) buffer (Sigma-Aldrich, R0278) supplemented with protease inhibitor (Roche, 11697498001) and maintained with constant agitation for 30 min at 4°C. Then the suspension was centrifuged at 4°C for 20 minutes at 12,000 rpm. The supernatant was collected as protein extract and quantified using BCA Protein Assay (Thermo Scientific, 23228).

Immunoblotting: 30 µg of the extracted protein in RIPA buffer were mixed with 1x reducing (10% b-mercaptoethanol) NuPAGE loading buffer (Life Technologies, Carlsbad, CA). After denaturation at 70°C for 10 min. The protein was loaded on a precast polyacrylamide NuPage 4-12% Bis-Tris protein gel (Invitrogen, Carlsbad, CA, USA) and 1x MOPS (1M MOPS, 1M TrisBase, 69.3mM SDS, 20.5mM EDTA) to be separated according to the length using SDS -PAGE (100V, 200mA, 2h). Proteins were blotted on 0.45 µm pore size Immobilon-P Polyvinylidene difluoride (PVDF) membrane (EMD Millipore, Billerica, MA; USA). The membrane was pre-activated for 20 sec in methanol and equilibrated in 1X NuPage Transfer buffer (1.25 mM bicine, 1.25 mM BisTris, 0.05 mM EDTA, and 10% ethanol) for 30 min at RT. The membrane was blocked in 5% bovine serum albumin for 1 h at RT and incubated overnight at 4°C with primary antibodies: Anti-CEP83 produced in rabbit (1:500, Sigma-Aldrich) and Anti-α-Tubulin produced in mouse (1:500, T9026, Sigma-Aldrich). The membrane was incubated for 1 h at RT with horseradish peroxidase-conjugated secondary antibodies (Sigma-Aldrich, Saint Louis, MO, USA) with 1:2000 dilution. Chemiluminescent reagent (Super Signal–West Pico; Thermo Scientific, Waltham, MA; USA) was used to detect the proteins. The spectra^TM^ Multicolor Broad Range Protein Ladder (Thermo Fisher Scientific, USA) was used to evaluate the molecular weight of corresponding protein bands.

### Histology and Immunofluorescence (IF) staining

After organoid fixation in BD Cytofix buffer (554655, BD Biosciences) for 1 hour on ice, the organoids were gradually dehydrated in increasing ethanol concentrations for 15 minutes each. Then organoids were cleared in xylene for three times 20 minutes each. After infiltration with melted paraffin at 65°C three times for 30 minutes each, the organoids were embedded in paraffin and processed in 3.5 µm-thick sections using a HM 355S microtome. The sections were deparaffinized, dehydrated, and stained with hematoxylin (Sigma-Aldrich, Saint Louis, MO) for 3 minutes and in 1% eosin (Sigma-Aldrich) for 2 minutes. The sections were mounted using Kaiser’s glycerol gelatin-based mounting medium. Images were captured with a Leica CTR 6000 microscope (Leica Biosystems, Wetzlar, Germany).

For immunostsining, cultured cells (D7) and organoids (D25) were fixed with BD Cytofix for 10 miuntes on ice. Then cells were permeabilized with BD Perm/Wash (554723, BD Biosciences), twice, 15 minutes per each. Cells were blocked with blocking solution (1% BSA + 0.3% triton-X-100 in 1X DPBS) for 2 hours at RT or overnight at 4°C. Cells were incubated overnight at 4°C with primary antibodies (table S2). Cells were then washed twice (10 min each) and incubated with fluorescence-labeled secondary antibodies with 1:500 dilution including Cy3, Cy5, Alexa488, and Alexa647 (Jackson ImmunoResearch, Newmarket, UK) and Cy3 Streptavidin (Vector lab, Burlingame, USA) overnight at 4°C. DAPI was then used for nuclear staining (Cell signaling Technology, Danvers, MA, USA) with 1:300000 dilution for 1 hour at RT. Finally, cells were mounted with Dako fluorescent mounting medium (Agilent Technologies). The images were taken using a SP8 confocal microscope (Carl Zeiss GmbH, Oberkochen, Germany). All the quantitative analyses of the taken images were performed using ImageJ (1.48v; National Institutes of Health, Bethesda, MD) software.

### Supplemental Tables

**Table S1:**
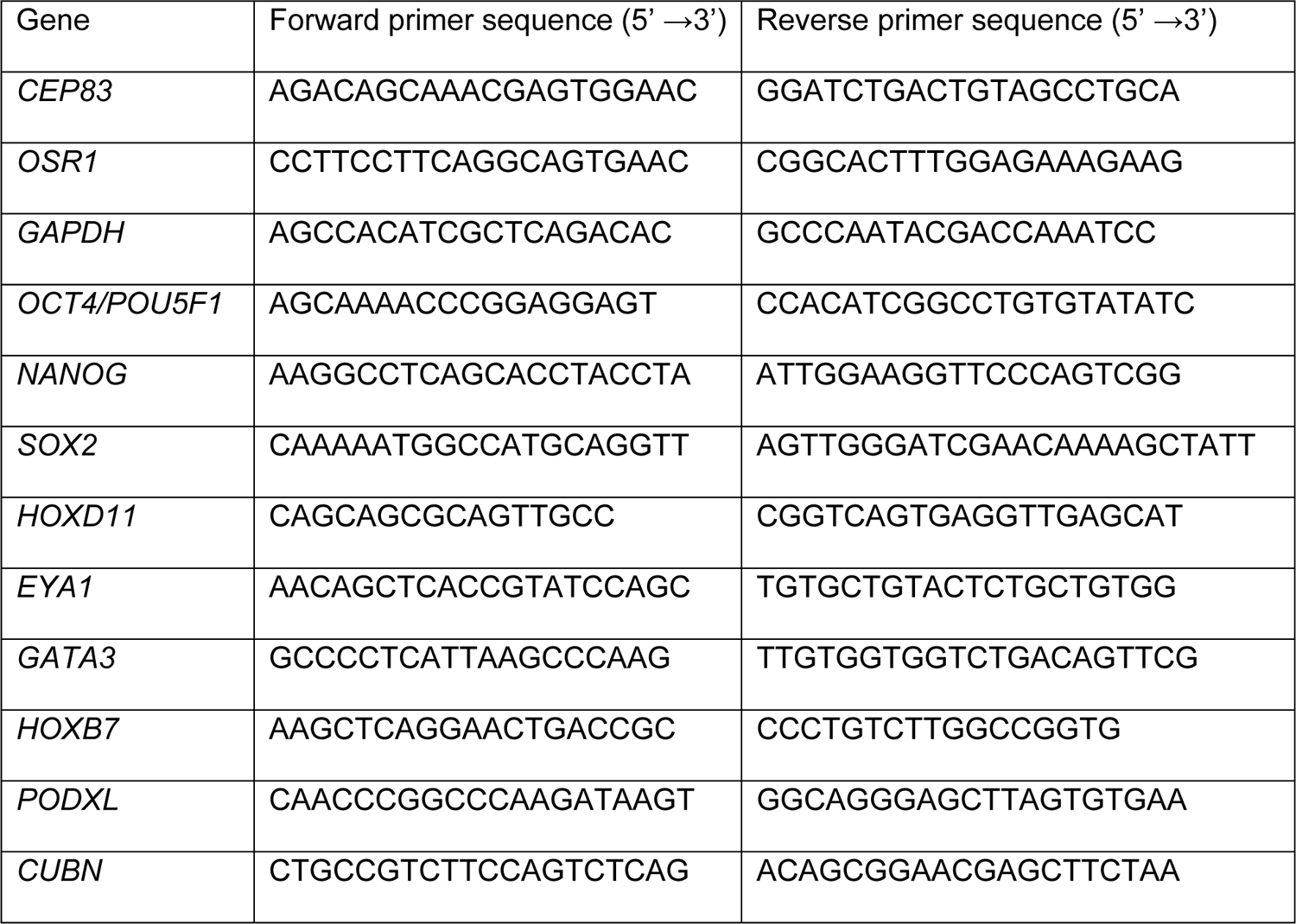
Primers list used in the qPCR.

**Table S2:** Primary antibodies used in IF staining.

| Antibody | Dilution | Company |
| --- | --- | --- |
| Monoclonal Anti-Tubulin, acetylated antibody (T6793). mouse | IF: 1:2000 | Sigma-Aldrich, Saint Louis, MO, USA |
| anti-Cdh1 | IF: 1:200 | BD Bioscience, San Jose, CA, USA |
| Lotus tetragonolobus lectin (LTL) | IF: 1:200 | Vector lab, Burlingame, USA |
| Anti-NPHS1 | IF: 1:300 | R&D System, Minneapolis, MN, USA |
| Anti- CEP83 | IF: 1:200 | Sigma-Aldrich, Saint Louis, MO, USA |

